# Integration of multi-modal measurements identifies critical mechanisms of tuberculosis drug action

**DOI:** 10.1101/2024.11.20.623860

**Authors:** William C. Johnson, Ares Alivisatos, Trever C. Smith, Nhi Van, Vijay Soni, Joshua B. Wallach, Nicholas A. Clark, Timothy A. Fitzgerald, Joshua J. Whiteley, Artem Sokolov, D. Michael Ando, Dirk Schnappinger, Kyu Y. Rhee, Bree B. Aldridge

## Abstract

Treatments for tuberculosis remain lengthy, motivating a search for new drugs with novel mechanisms of action. However, it remains challenging to elucidate the direct targets of a drug, and even more so, to determine which disrupted cellular processes lead to bacterial killing. We developed a computational tool, DECIPHAER (DEcoding Cross-modal Information of PHarmacologies via AutoEncodeRs), to select the important correlated transcriptional and morphological responses of *Mycobacterium tuberculosis* to drug treatments. By finding a reduced feature space from these measurements, DECIPHAER highlighted essential features of Mtb cellular damage such as phosphosugar stress and inhibition of translation and DNA replication. After training, DECIPHAER provides cell-death-relevant insight into single-modal datasets, enabling interrogation of drug treatment responses for which transcriptional data are unavailable. Using morphological data alone with DECIPHAER, we discovered that respiration inhibition by the poly-pharmacological drugs, SQ109 and BM212, can influence cell death more than their effects on the cell wall. This study demonstrates that DECIPHAER can extract the critical shared information from multi-modal measurements to identify cell death-relevant mechanisms of TB drugs.

## INTRODUCTION

Tuberculosis (TB) is caused by *Mycobacterium tuberculosis* (Mtb) infection and killed ∼1.5 million people in 2021^1^. One reason for TB’s significant death toll is that treatments are too lengthy. The current treatment regimen involves a 4-6 month course of multiple antibiotics^1^. Substantial resources have been devoted to developing anti-TB drugs with new mechanisms of action (MOAs) to shorten the course that must be followed by patients^2^. MOA studies in TB rely heavily on characterizing the primary target of a drug by isolating and performing DNA sequencing on resistant mutants^3–6^. However, there are cases in which spontaneous resistance mutants against a compound are difficult to isolate, especially when the drug targets multiple cellular processes^7–9^. Moreover, primary target inhibition is not always the mechanism that causes cell death. In some cases, it is a downstream product or toxic byproduct of the targeted biological pathway that leads to bacterial killing^10–12^. To determine which drug-disrupted pathways lead to killing, we require MOA discovery methods that comprehensively measure the cellular processes downstream of primary target inhibition.

Unbiased omics approaches to studying TB drug MOA are promising as they offer a holistic readout on cellular state and can therefore capture pathway changes downstream of a drug’s primary target. Transcriptomic profiling allows simultaneous measurement of many cellular processes after Mtb is exposed to a drug. These analyses often focus on the most differentially expressed gene programs. However, because so many genes are differentially expressed in response to a drug treatment, a challenge has been defining the cell-death-relevant transcriptional response^13^. Another omics approach, morphological profiling, is effective at identifying the major disruptions in cellular processes that lead to Mtb deterioration^14,15^. By measuring changes in cell appearance, morphological profiling details the chemical and physical changes of cells after drug treatment. A shortcoming to this approach is that it does not provide a gene-, protein-, or molecule-level readout, making follow-up studies on MOA insights difficult.

We hypothesized that integrating the molecular detail of RNA-seq with the cell-damage-revealing information obtained by morphological profiling would complement each approach’s shortcomings and provide novel insight into drug mechanisms of action. Many multi-modal integration techniques have been developed to analyze data of similar structures, such as proteomics, chromatin accessibilities, and transcriptomics^16–21^. A common approach is to decompose each data modality into lower dimensional latent features that describe the most correlated multi-omics signatures. These latent features can then be used for biomarker discovery or analysis of cross-modal feature relationships. Many methods perform best when both omics measurements are made on the same population of cells (paired measurement)^22^, which is difficult with current bacterial omics technology. Additionally, these methods are most successful in cases where measurements from different modalities have similar features (proteins and transcripts can be mapped to the same gene). Specialized neural networks called autoencoders (Box 1) have recently proven powerful in these areas, as they can learn complex patterns and compute latent spaces from unpaired modality measurements with dissimilar features^23–25^.

We leveraged autoencoders to integrate RNA-seq and morphological profiling data in a shared latent space that is informative of TB drug MOA. Here, we describe this pipeline, called DEcoding Cross-modal Information of PHarmacologies via AutoEncodeRs (DECIPHAER). Our model enabled us to define a multi-modal latent space shared between unmatched samples of imaging and transcriptomics data. Using this aligned space, we identified correlated morphological and transcriptional features of cell damage, such as cell size, ribosome availability, and DNA replication and repair, consistent with previous studies in bacteria. We also identified a correlation between the TreS/GlgE pathway and nucleoid size, which was related to sugar-phosphate metabolism. Follow-up analysis of this pathway showed that sugar-phosphate poisoning might contribute to the mechanism of action of linezolid against Mtb. DECIPHAER was also capable of predicting multi-modal latent space- and transcriptional-signatures of TB drugs from morphological information alone. We used this function to investigate an MmpL3-inhibitor for which we had only measured morphological data: SQ109. While single-modal analysis revealed only the cell wall-damage effect of SQ109, analysis of the DECIPHAER multi-modal latent space highlighted its impact on cellular respiration. We confirmed that SQ109’s respiration effect impacts cell death more than its cell wall effect in some conditions, and our data suggests that this may be common among other MmpL3 inhibitors. This work establishes a quantitative framework for investigating the mechanisms of cell death by TB drugs and demonstrates the advantage of multi-modal analysis for highlighting critical features not obvious in single-modal analyses.

## RESULTS

### DECIPHAER integrates and translates distinct modality measurements of Mtb drug response

To generate comprehensive drug-response profiles for multi-modal analysis of MOA, we treated Mtb with 17 drugs that are in use or in development for TB treatment. In this drug set, mechanisms of action span multiple cellular targets, including respiration, cell wall synthesis, protein synthesis, etc. (Supplementary Table 1). We measured gene expression using RNA-seq (Methods) after both 4 hours and 24 hours of drug treatment to capture both early- and delayed-drug-action transcriptional responses. We generated a complementary morphological profiling dataset using a set of 14 drugs that overlap with our RNA-seq dataset. To capture the most dynamic morphological responses, Mtb were fixed after 17 hours of drug treatment^15^ (Figure 1A, left). We stained Mtb with membrane and nucleoid dyes and extracted single-cell morphological features such as staining intensity, cell area, and other mathematical descriptors of shape (Methods).

**Figure 1.**
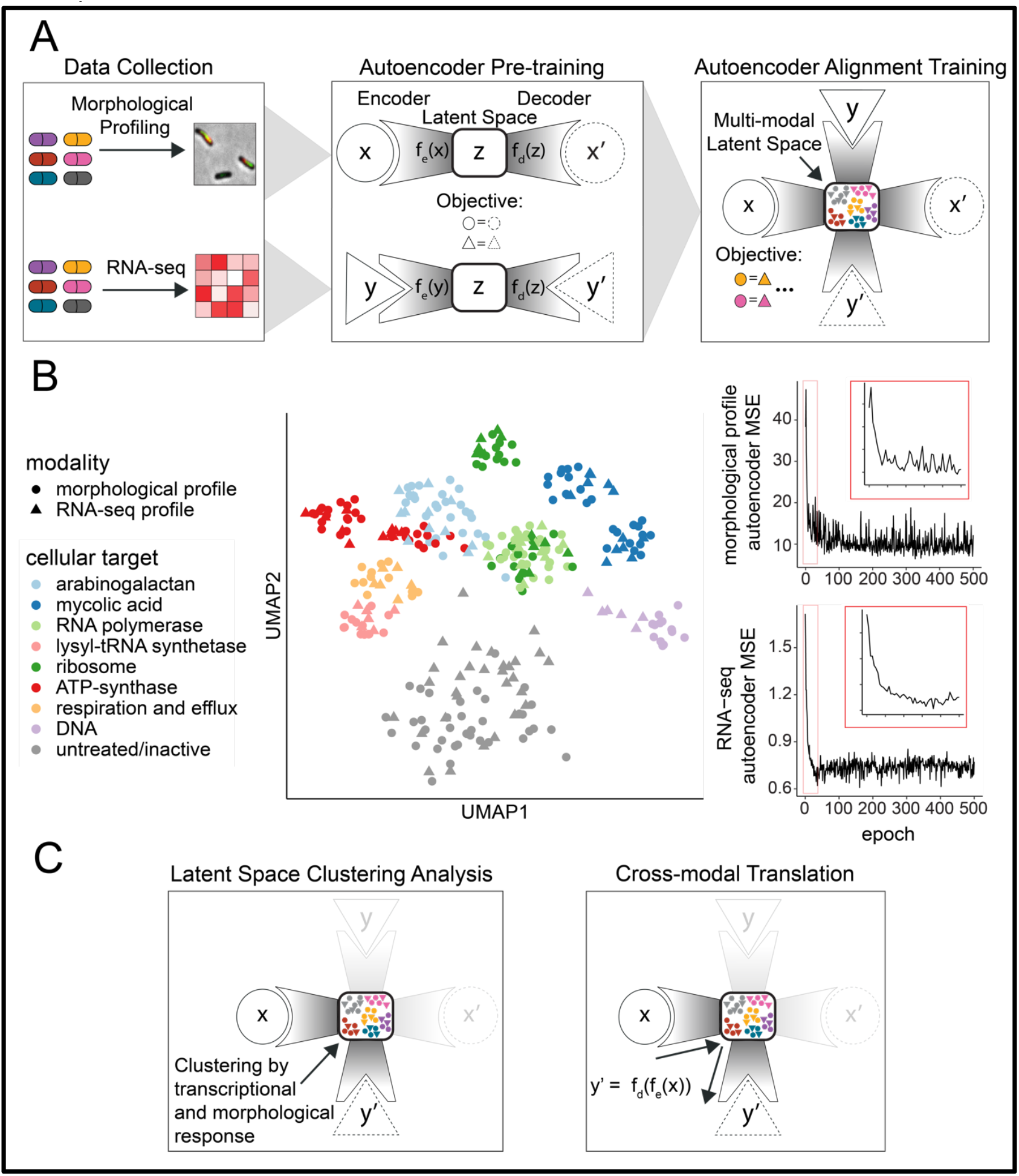
DECIPHAER computational framework integrates multi-modal measurements in a shared latent space and enables cross-modal prediction. (A) (Left) Depiction of drug mechanism of action profiling. Mtb cultures were exposed to a set of drugs spanning diverse mechanisms of action (colors), and the morphological and transcriptional response of the bacteria were measured. (Center) Single-modal autoencoders are trained on RNA-seq (triangles) and morphological profiles (circles). By optimizing reconstruction from the latent space bottleneck (objective), the autoencoders learn modality-specific low-dimensional latent features (Z or Latent Space). (Right) DECIPHAER alignment training adjusts the parameters of the pre-trained modality-specific autoencoders to align the two modalities’ latent feature distributions. The training objective ensures morphological and transcriptional data of similar drug-response phenotypes are embedded in similar latent space locations. By doing this, the multi-modal latent space is optimized to contain only the most correlated morphological and transcriptional features, while the information with low correlation across the modalities is excluded. (B) (Left) UMAP of the 50-dimensional DECIPHAER multi-modal latent space. Data point colors indicate the drug’s primary target and the shape indicates the modality (RNA sample from a specific time point and biological replicate, triangle; morphological profile summarized from 1,000 randomly sampled single cell profiles, circle). (Right) Mean-squared error loss (MSE) (y-axis) as a function of training time, in epochs (x-axis), from the morphological profile (top) and RNA-seq (bottom) autoencoders trained on a shared latent space. (C) Schematic of the analyses that can be performed using the DECIPHAER model. (Left) The multi-modal latent space permits clustering and phenotype analysis of drug mechanisms based on the correlated morphological and transcriptional drug response of Mtb. (Right) Matching one modality’s encoder with another’s decoder creates a cross-modal translation model, enabling the prediction of modality y from modality x.

To integrate the two data modalities, we aimed to calculate a space that would capture only the transcriptional features with the highest correlation with our morphological dataset while omitting features in each dataset with distinct information. We hypothesized that by reducing the dimensionality of these two datasets to a set of variables that robustly summarizes each, we could better explain TB drug MOA. We trained autoencoders (Box 1; Figure 1A, center) to compress our two datasets into latent features, which are mathematical composites of the most correlated transcriptional and morphological features. By optimizing a classifier on the latent space that ensures similar drug phenotypes (i.e., measurements from drugs with similar mechanisms) form clusters, regardless of the modality from which they came, the latent space omits modality-specific information but retains relevant biological information (Figure 1A, right and Methods). Unlike most other methods, this framework does not require paired measurements of gene expression and morphology to perform training and integration, allowing us to use the latent space to explore Mtb’s response to drugs for which we do not have both morphology and gene expression measurements (Supplementary Figure 1). Training of the autoencoders to embed RNA-seq and morphological data into the same latent space resulted in the convergence of the reconstruction loss (mean-squared error) of each modality (Figure 1B, right). This reduction in model error suggests that there is significant mutual information between the modalities since each dataset can be accurately reconstructed from the same latent space.

If the autoencoders’ latent space holds a joint representation of transcriptional and morphological data that describes the response of Mtb to drug treatment, then this space should discriminate cells treated with drugs that induce disparate responses, and cluster ones that induce similar responses (Figure 1C, left). A uniform manifold approximation (UMAP) of DECIPHAER’s 50-dimensional latent space revealed clustering of morphological and RNA-seq data points by the drug treatments’ primary cellular targets, indicating an aligned and cell-response-informative latent space (Figure 1B, left). To quantify the degree of integration and the generalizability of the networks to new observations of RNA-seq and morphological data, we performed a k-nearest neighbors (kNN) analysis on the latent space. Using 6-fold cross-validation, we assessed the autoencoder’s accuracy at embedding morphological profiles near RNA-seq profiles of the same drug phenotype cluster and vice versa (Supplementary Figure 2A and Methods). The cross-modal autoencoders correctly embedded test morphological data points next to RNA-seq data points of the same drug phenotype cluster with a median accuracy of 93% (Supplementary Figure 2B). The cross-modal autoencoders did not generalize as well to new RNA-seq data points as the model obtained a median 59% kNN accuracy (Supplementary Figure 2C). This is likely because our held-out RNA-seq data contains RNA samples from Mtb treated with drugs for multiple durations, each of which have significant transcriptional differences to which our model cannot generalize (Supplementary Figure 3). To quantify how tightly data points of the same drug phenotype cluster together in the latent space we calculated an average F-test statistic (Methods) and found values significantly greater than 1 at a range of model hyperparameters (Supplementary Table 4). Together, these analyses suggest that our model can generate a latent space that discriminates drug phenotypes and generalizes well to morphological profiling test data.

Another use of our model is the ability to translate across modalities, enabling the prediction of corresponding gene expression changes given a morphological profile. This allows us to interpret how features in one modality relate to changes in the other and make cross-modal predictions from one modality alone. Because gene expression changes are more interpretable than morphological changes, we focused on the evaluation of our morphological profile encoder and RNA-seq decoder (Figure 1C, right). Translation of morphological profile test data to transcriptomics data revealed clustering of the predicted data with true transcriptomic profiles of the same drug phenotype cluster (Figure 2A). To quantify the accuracy of our model’s translations, we obtained sets of differentially expressed genes for each drug treatment in our RNA-seq dataset (Methods). For each drug’s gene set, we calculated a Pearson’s correlation coefficient between the true transcriptional levels and the transcriptional levels predicted by new morphological data with 6-fold cross-validation (Figure 2B). We observed a strong correlation between predicted and true transcriptional levels for the same drug (mean R = 0.73) and weak correlation between different drugs (mean R = -0.07) (Figure 2B). The predicted transcriptional profiles for rifampicin (RIF), rifapentine (RFP), and linezolid (LIN) correlated strongly with the true transcriptional data for linezolid (mean R = 0.86, 0.95, and 0.87 respectively), suggesting a highly similar phenotype between the three drug responses. Because rifampicin and rifapentine primarily inhibit RNA polymerase^26,27^ and linezolid inhibits the ribosome^28^, the correlation between their predicted transcriptional data may be explained by shared effects on protein production. These data demonstrate that our cross-modal autoencoders can accurately predict transcriptional data relevant to drug mechanism of action, given only morphological profiles.

**Figure 2.**
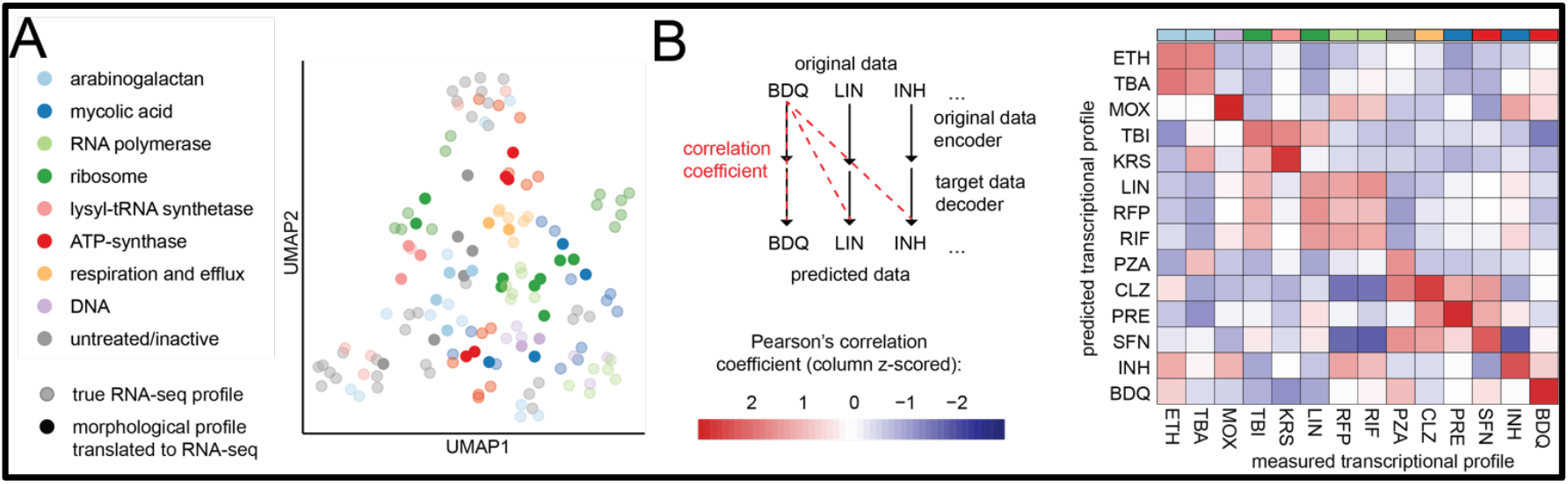
Computational validation of DECIPHAER to ensure consistent alignment and accurate cross-modal predictions. (A) Representative UMAP of a single test split of morphological profiling data translated to RNA-seq with true RNA-seq plotted. The transparency indicates the modality, and the colors indicate the cellular target of the drug treatment. Each translated data point (solid) represents the transcriptomic profile predicted from a test morphological profile (summarized from 1,000 randomly sampled single cell profiles) treated with a drug (color). Each true RNA-seq data point (transparent) represents a training transcriptomics profile from a biological replicate of a drug treatment at the 4-or 24-hour timepoint. (B) Approach (left) and quantification (right) to assess cross-modal prediction accuracy. Genesets were calculated for each drug treatment (Methods), and each compound’s held-out morphological profile test data was translated into a transcriptomic profile. The Pearson’s correlation coefficient between the true RNA-seq and the predicted RNA-seq is calculated among the genes from each drug treatment’s geneset. Accurate predictions should show strong correlations between translated and true RNA-seq data of the same drug but low correlations between data from different drugs. The mean Pearson’s correlation coefficients of DECIPHAER cross-modal predictions from 6-fold cross-validation test data are plotted in heatmap form (right). True RNA-seq are in columns and the RNA-seq predicted from morphological data in rows. Columns were converted to z-scores to show relative differences between predictions (red = high correlation, blue = low correlation). Column colors indicate the cellular target of the drug treatment.

### Box 1

Dimensionality reduction involves reducing the number of features in a dataset while retaining its relevant information. This reduced feature space, often referred to as a latent space, can provide meaningful insight on an otherwise convoluted dataset. Autoencoders are well-suited to the task of dimensionality reduction. These neural networks consist of two modules: an encoder and a decoder. The encoder network is trained to compress input data into a lower dimensional latent space. The decoder network is optimized to take the data encoded in this latent space and transform it to match the input information. The latent space learned thus reduces the many features of the dataset into a few latent dimensions which maintain maximal information (Figure 1A, center).

To generate latent features that are composites of more than one modality, i.e., morphological or transcriptomics datasets, the objective used to train the model must be augmented. For this task, the latent features must maximize the information from all modalities, and to ensure consistent alignment, they should cluster multiple modality observations (RNA samples, morphological profiles) from the same biological group (drug-treatment, drug-induced phenotype etc.) together (Figure 1A, right, colors). Such a model can be difficult to fit to data where paired-measurements, or an empirical mapping of an observation in modality *x* to the same observation in modality *y*, are not feasible to measure, as is the case for our morphological and transcriptional datasets. In this work, we approximated this mapping using a framework that relies on aligning the distributions of similar biological groups from the two modalities. To perform this alignment, we utilized the loss functions proposed by Yang & Belyvea *et al*.^23^ to train a classifier on the latent space which minimizes the distance between data points of the same biological group, regardless of the modality from which they came. Through this method, the latent space learned by the autoencoders omits modality-specific features and retains relevant biological information.

#### Feature

variable that measures some quantifiable attribute. A feature in an RNA-seq dataset would correspond to an individual gene’s expression level whereas a feature in a morphological dataset would correspond to a cell appearance descriptor, like cell area. Feature values are measured across different observations, ie. cells, biological replicates.

#### Latent space

low-dimensional feature space that summarizes the major patterns that exist in the original higher dimensional dataset.

#### Encoder

neural network trained to compress high dimensional input data into a lower dimensional representation.

#### Decoder

neural network trained to reconstruct true data from the latent space representation of the data.

#### Objective/loss

function that quantifies the accuracy of a model. The model is trained to either maximize or minimize the value of the objective/loss function.

#### MSE (mean-squared error)

the average difference between the true values and the values predicted by a model. A small MSE value indicates the model is accurate.

### DECIPHAER highlights correlated transcriptional and morphological features of Mtb damage

One major goal in multi-omics analyses is to identify features with mutual information across different omics modalities, as these will be robust signatures that are invariant to unwanted artifacts from measurement types. In our application, we aimed to determine the most robust signatures of drug-induced cellular damage in Mtb; therefore, we sought to identify the most correlated morphological and transcriptional features. To calculate correlation coefficients among features from different data modalities, paired measurement of gene expression and morphology is usually required, which was not feasible in our study. Instead, our trained autoencoders can be used to generate predictions of paired omics measurements (Supplementary Figure 4A). Using this methodology, we calculated all pairwise cross-modal feature correlation coefficients. Many feature pairs were predicted to be strongly correlated, with 18,819 cross-modal feature pairs having Spearman rank correlation coefficients greater than an absolute value of 0.5 (Supplementary Figure 4B).

To determine the characteristic biological pathway changes associated with each Mtb morphological feature, we performed pathway enrichment analysis on the most correlated gene transcripts (Methods). Cell size features were enriched for ribosome biogenesis and DNA replication and repair gene expression programs (Figure 3A, center panel). Ranking cells according to their area and translating to predicted gene expression levels demonstrated that increases in cell size corresponded to the upregulation of genes involved in DNA replication and repair and ribosome biogenesis (Figure 3B). This relationship was a common damage response to drug treatments that target DNA and protein synthesis; concordantly, drugs targeting either protein synthesis or DNA caused increases in Mtb cell area (Supplementary Figure 5).

**Figure 3.**
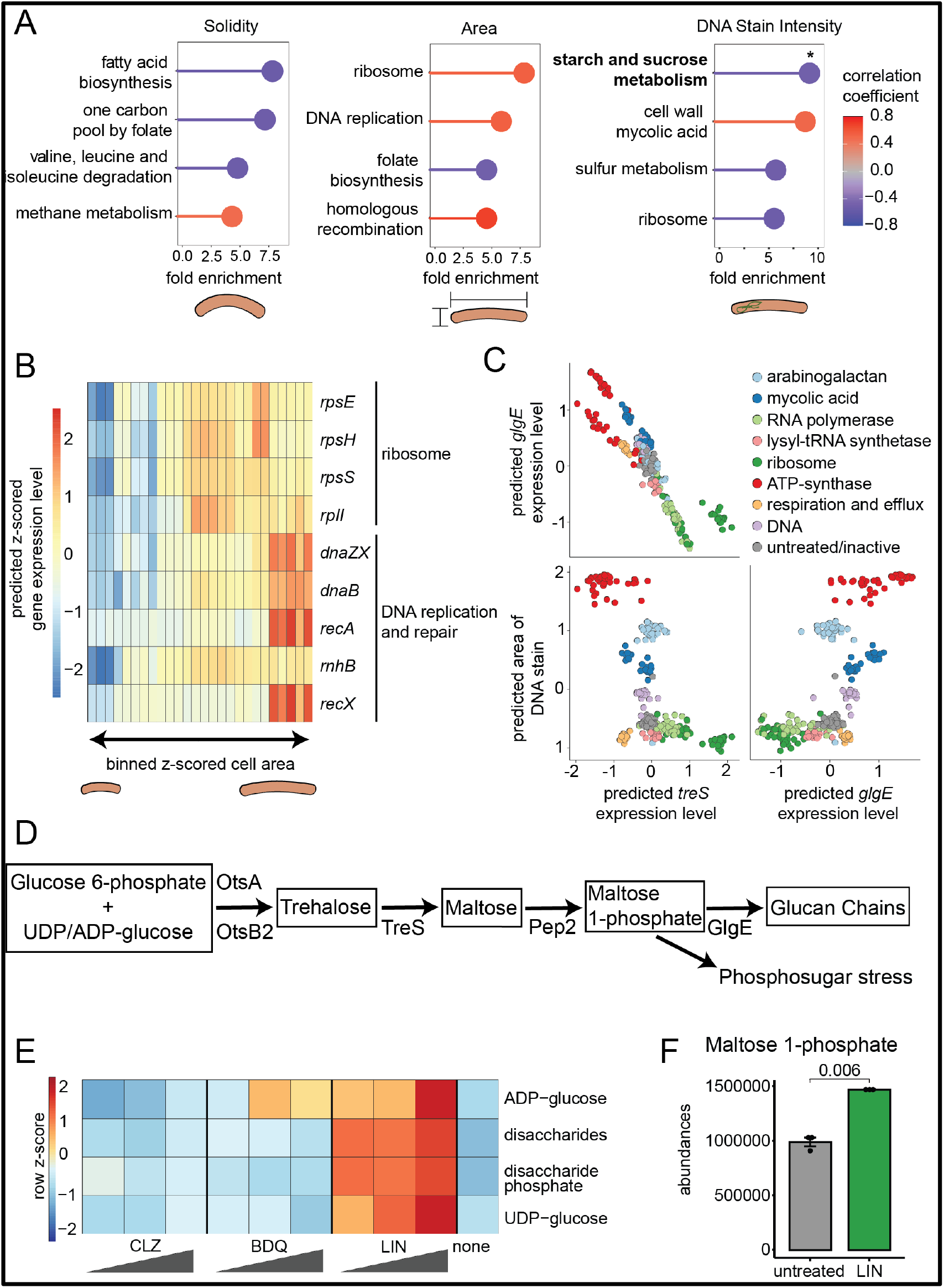
DECIPHAER identifies correlated gene expression and morphology relationships that are informative of drug mechanism. (A) Pathway enrichment ratios (PathFindR) of three morphological features. For each morphological feature, the most correlated gene expression features (Spearman’s rank correlation coefficient > 0.5 or < -0.5) were selected for pathway enrichment. Top enriched pathways are colored by their average Spearman rank correlation coefficient and the given morphological feature is shown below (positive correlation, red; negative correlation, blue). From left to right, the morphological features are solidity, cell area and DNA staining intensity (the * indicates the sugar metabolism pathway of which TreS and GlgE are members). (B) Heatmap of the predicted expression levels for genes involved in ribosome biogenesis and DNA replication and repair across the entire range of possible cell area bins. (C) Scatter plot of DECIPHAER-paired morphological and RNA-seq data for *treS, glgE* and area of nucleoid stain. Each feature (axis) is converted to a z-score to show relative differences between drug treatments. Each data point represents either a morphological profile summary of 1,000 cells and its corresponding DECIPHAER-predicted gene expression level or an RNA sample and its corresponding DECIPHAER-predicted nucleoid staining area. (D) Metabolic map of the maltose 1-phosphate pathway depicting accumulation of sugar-phosphate in linezolid treated Mtb. (E) Abundances of metabolites for the KEGG maltose-1-phosphate pathway in cholesterol-adapted Mtb cultures treated with drugs for 24 hours. Rows are z-scored to demonstrate relative levels of the metabolites. Bedaquiline and clofazamine are shown as control drug treatments. Three doses are shown for each compound at 1x, 5x and 10x their IC50 (n = 3, 2 independent experiments). (F) HILIC-MS quantification of maltose-1-phosphate in cholesterol-adapted 24-hour-linezolid-treated and untreated Mtb cultures (p-value of a two-sided, unpaired t-test, n = 3).

The correlation between cell area, translation, and DNA homeostasis was consistent with observations in the related organism, *Mycobacterium smegmatis* (*M. smegmatis*)^29,30^. Inhibiting DNA replication in *M. smegmatis* limits the amount of genetic material required for cells to divide, leading to elongated cells. The mechanism behind the relationship between cell area and translation inhibition in *M. smegmatis* is less understood. In *E. coli*, cell size and protein translation correlate with the cell cycle^31,32^. Therefore, the increased cell area in Mtb caused by protein synthesis inhibitors may be due to an effect on the cell cycle. Together, these data suggest that DECIPHAER can capture gene-morphology relationships in drug perturbation data consistent with previous observations in related organisms. It also indicates that drug perturbations to DNA and translation induce the strongest transcriptional and morphological damage responses in Mtb.

### Linezolid treatment alters phosphosugar metabolism and cell staining features

One application of DECIPHAER is to determine correlations between morphological and gene expression measurements to identify strong signatures of drug-induced cellular damage. One of the gene expression pathways with the most mutual information with our morphological profiling dataset was the starch and sucrose metabolism pathway, GlgE/TreS. This pathway was highly correlated with many cell staining features, notably DNA staining intensity (Figure 3A, right, star; and Supplementary Figure 6). TreS catalyzes the conversion of trehalose to maltose and GlgE utilizes the phosphorylated maltose substrate, maltose 1-phosphate (M1P), to generate α-1,4-glucan chains (Figure 3D)^33^. *glgE* is essential because its deletion reduces the incorporation of M1P into glucan chains, leading to a toxic build-up of free M1P and cell death.

Because GlgE- and TreS-mediated sugar metabolism influences Mtb viability, we sought to determine which drug treatments in our dataset effect this pathway to the greatest effect. We plotted the paired omics measurements predicted by DECIPHAER and determined that Mtb treated with 50S ribosomal subunit inhibitors caused decreased *glgE* expression and DNA stain area and a corresponding increase in *treS* expression (Figure 3C, green points). This prediction was consistent with measured RNA-seq data, in which the 50S ribosomal subunit inhibitor, linezolid, caused the greatest decrease in *glgE* expression among our drug-set (Supplementary Figure 7A). Because GlgE prevents M1P build-up, we hypothesized that the linezolid-induced downregulation of *glgE* expression could cause phosphosugar stress. In support of this, an RNA-seq analysis of linezolid-treated cells identified significant remodeling of components involved in respiration, a known response to phosphosugar stress^33^. Specifically, linezolid treatment caused reduced expression of ATP synthase and cytochrome *c* oxidase components (Supplementary Figure 7B)^33^.

To evaluate whether linezolid-treated cells experience high levels of sugar phosphates, we performed LC-MS on Mtb treated with linezolid and analyzed the metabolic changes within the TreS/GlgE pathway. A dose-dependent accumulation of nucleotide sugars (which can be converted into M1P (Figure 3D)), disaccharides, and, importantly, disaccharide phosphates, of which M1P is a species, further supported the hypothesis that sugar-phosphate poisoning may be a component of linezolid’s mechanism (Figure 3E). Bedaquiline and clofazimine treatments were used as controls because they did not induce downregulation of *glgE* expression (Supplementary Figure 7A), and, as such, did not cause sugar phosphate accumulation (Figure 3E). Using HILIC-MS (Methods), we determined that linezolid does indeed increase M1P levels (Figure 3F). Together, these data demonstrate that linezolid treatment induces elevated sugar-phosphate levels and a sugar-phosphate stress response, suggesting phosphosugar poisoning may contribute to the potency of linezolid against Mtb.

### Single-modal data can be evaluated using DECIPHAER’s latent- and cross-modal predictions

Predicting multi-modal data from single-modal measurements is of great interest to biologists because performing multiple omics assays is often prohibitively time and resource-intensive. Perhaps just as useful is a tool that can identify the “important” features of an omics dataset based on those features’ correlation with another data modality. The ability of DECIPHAER to accurately translate independent test splits of morphological profiling data to latent space and RNA-seq feature values (Figure 2A/C and Supplementary Figure 2) raises the possibility that the model may also be useful for analyzing drug treatment profiles for which we have only collected morphological data.

To determine whether we could predict latent space features of compounds for which we had only measured morphological data, we encoded morphological profiles from a set of 34 compounds for which no RNA-seq had been collected into the DECIPHAER latent space (Supplementary Table 1). Then, we performed hierarchical clustering on these compounds’ latent embeddings along with all training data to determine which training-set drug treatment responses were most similar (Methods). Of the active drugs (Methods) for which we only measured morphological profiling data, 68% shared their most recent clade with training drugs of the same expected broad cellular target (Supplementary Table 5). This suggests that the DECIPHAER multi-modal latent space can be used to investigate the mechanism of action of drugs for which only one modality measurement has been collected.

We then sought to assess whether DECIPHAER’s translator function could predict transcriptional changes from the morphological data of active drugs for which we did not measure RNA-seq. We decoded the latent embedded morphological profiling data from these drugs to RNA-seq and quantified the Jaccard similarity coefficient between their differential expression profiles and those of the training drugs (Methods). By this metric, the broad cellular target of the most similar training drugs’ differential expression profile matched the expected broad cellular target of the drug 65% of the time (excluding drugs that did not reach IC50 in the standard growth condition) (Supplementary Table 6).

To gain further confidence in the DECIPHAER predictions from morphological data alone, we evaluated three use cases in depth, focusing on drugs that are known to target distinct cellular processes: DNA replication, cellular respiration, and RNA polymerization. Morphological profiles of the DNA-acting drugs, mitomycin, levofloxacin and daunorubicin^34–36^, clustered with the DNA-acting training drug, moxifloxacin, by UMAP and hierarchical clustering (Figure 4A). We used DECIPHAER’s translator function to predict transcriptional changes from these morphological profiles and found that their differential expression profile was most similar to that of moxifloxacin (Figure 4B). Pathway enrichment analysis demonstrated that genes involved in homologous recombination, mismatch repair, and DNA replication were predicted to be significantly upregulated in response to these drug treatments, a response consistent with previous transcriptional analysis on quinolone-treated Mtb^13,37^ (Figure 4C/D).

**Figure 4.**
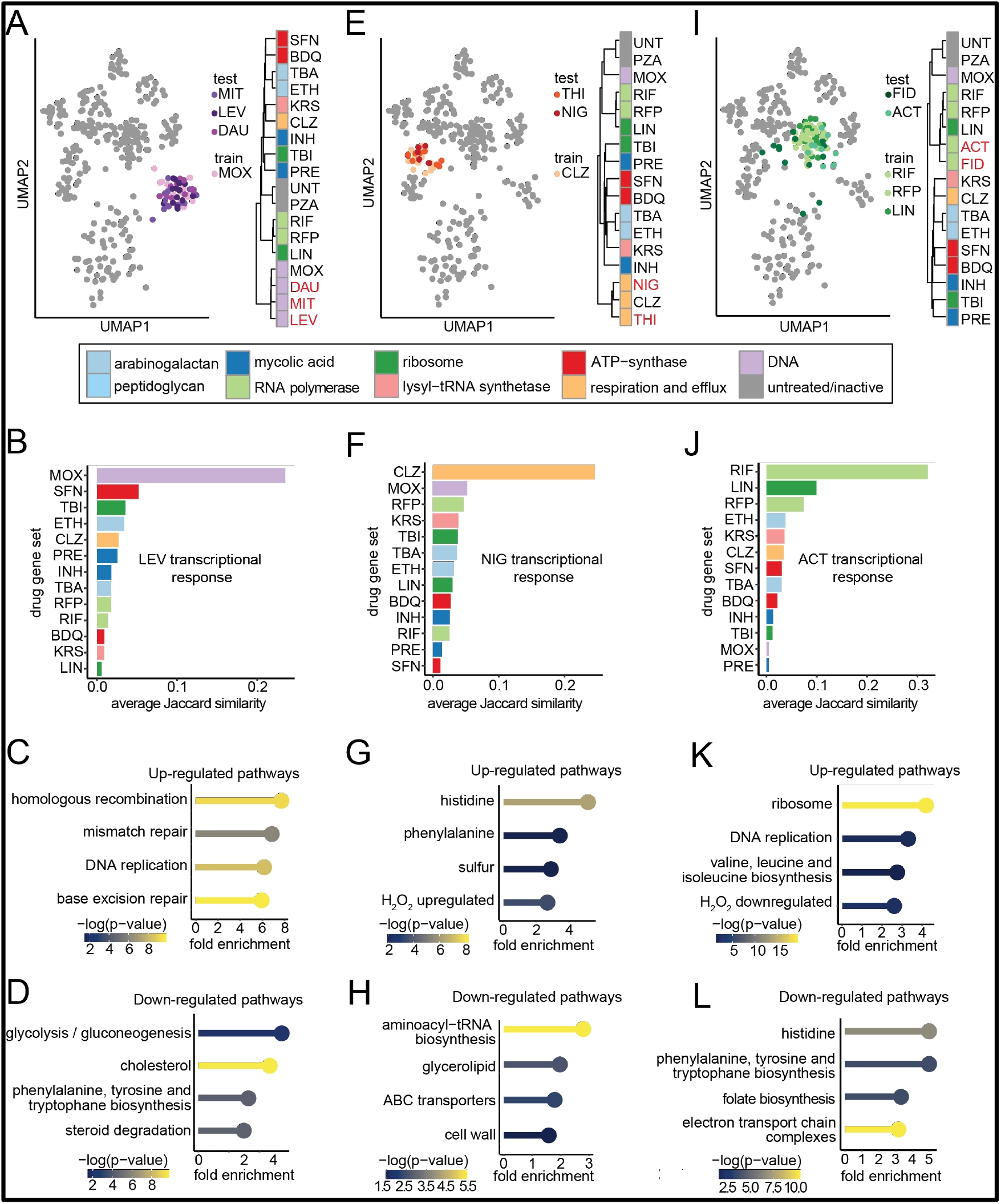
DECIPHAER predicts latent features and transcriptional responses from morphological measurements of Mtb treated with model-unseen drugs. (A, E, I) UMAP plots (left) and dendrograms (right) of DECIPHAER latent embedded morphological profiles for which there was no available RNA-seq data (test and red in the dendrogram), along with the training compounds’ latent embeddings (train). The hierarchical clustering was performed on the median latent feature values for each drug using Manhattan distance and Ward2 clustering. Colors indicate the cellular target of the drug. (B,F,J) Average Jaccard similarity coefficient between morphological data of drug morphological data translated to RNA-seq and all training-set RNA-seq (Methods). (C,D,G,H,K,L) Up- and down-regulated pathway enrichments from RNA-seq predicted from morphological data. Differential expression testing for enrichment analysis in C,D; G,H; and K,L; was performed on the translated data from the clusters in A, E and I, respectively, against all other training-set translated RNA-seq using Wilcoxon-rank sums. The color of the bubble plot indicates the highest -log_10_(p-value) among the differentially expressed genes in the pathway and the x-axis indicates the fold enrichment of the pathway (PathFindR).

Nigericin and thioridazine, which target respiration through membrane potential dissipation and electron transport chain (ETC) inhibition^38–40^, respectively, were most similar to the ETC targeting drug, clofazimine in the latent and gene expression spaces (Figure 4E & 4F). Thioridazine and nigericin were predicted to cause upregulation of the H_2_O_2_-responsive gene cluster, indicating redox homeostasis disruption (Figure 4G)^13^. They were predicted to cause significant downregulation of aminoacyl-tRNA biosynthesis which is known to occur in response to respiration-targeting drugs, likely because of declining numbers of charged tRNAs (Figure 4H)^13^. Fidaxomicin and actinomycin D inhibit transcription by interfering with RNA polymerase to DNA binding^41,42^. In the latent and predicted gene expression spaces, these two drugs were most similar to drugs targeting RNA-polymerase, rifampicin and rifapentine, as well as the protein synthesis inhibitor, linezolid (Figure 4I & 4J). Pathway enrichment analysis of Fidaxomicin and actinomycin D indicated a response indicative of RNA and protein synthesis cessation, with ribosome biogenesis being the most enriched pathway among all up- and down-regulated genes (>4-fold enrichment and -log_10_(p-value) > 15) (Figure 4K & 4L).

Together, this unsupervised analysis demonstrates that the DECIPHAER latent space features can be used to investigate mechanisms of drugs for which we only have morphological data. Additionally, it demonstrates that the RNA-seq predictions made by our model given only morphological profiling data of drugs align with what is expected of their purported mechanism of action.

### The multi-modal latent space highlights the respiration effect of SQ109

In recent decades, there have been significant efforts to develop drugs with novel mechanisms against Mtb^2^. One particularly potent series that resulted from this work was the MmpL3 inhibitors^2^. These drugs interfere with the MmpL3 transporter involved in transferring mycolates to the cell wall core and thus target the mycolic acid layer of the cell wall^7^. For some of these compounds, secondary or downstream effects on the proton motive force (PMF) and respiration have also been observed, suggesting that they may interfere with ATP synthesis^43^. We sought to explore the mechanism of action of one such compound, SQ109, using the DECIPHAER framework. This analysis aimed to determine whether our model can highlight multiple mechanisms of action of compounds with poly-pharmacology and to determine the dominant mechanism of action of MmpL3 inhibitors.

To characterize the multi-modal phenotype of SQ109, we performed morphological profiling on Mtb treated with the drug and encoded this information into DECIPHAER. A UMAP of the DECIPHAER latent-embedded SQ109 morphological data revealed a lack of tight clustering with other training compounds, suggesting a mechanism of action distinct from the other cell wall-acting agents in our dataset (Figure 5A). Hierarchical clustering on the latent embeddings identified a clade containing SQ109 and the ATP-synthase inhibitors bedaquiline (BDQ) and TBAJ-876 (SFN) (Figure 5B), suggesting that the correlated morphological and transcriptional response of Mtb to SQ109 is most similar to respiration inhibitors. In contrast, a single-modal analysis of SQ109’s morphological and predicted transcriptional data determined its profile was most similar to other cell wall-acting agents (Supplementary Figure 8A & 8B).

**Figure 5.**
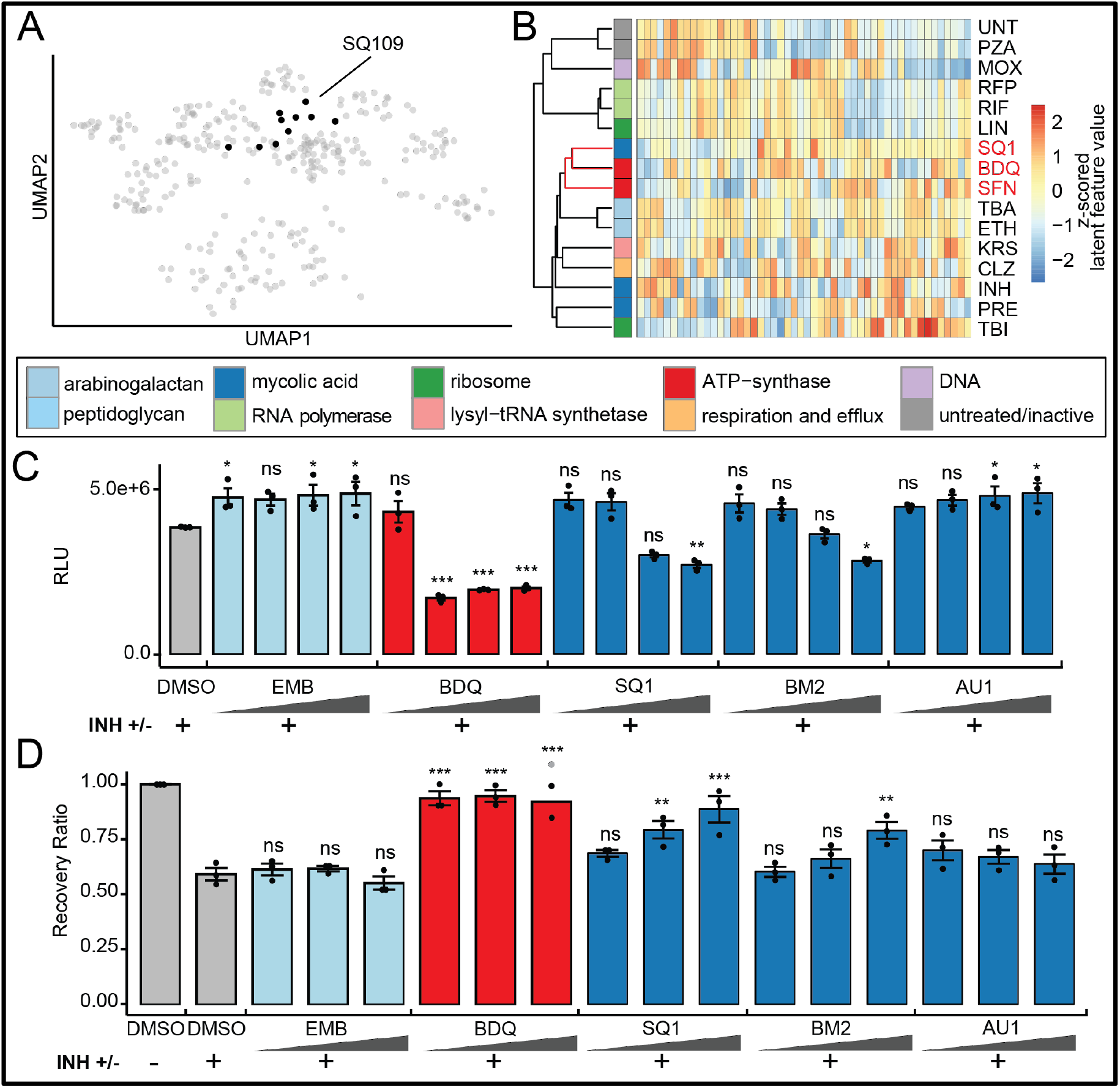
DECIPHAER’s latent space provides novel insights on the mechanism of action of SQ109. (A) UMAP plot of the DECIPHAER latent embedded morphological profiles of SQ109 (black), along with the alignment training compounds (gray). (B) Hierarchically clustered heatmap of the medians of SQ109’s and the training-set drugs’ DECIPHAER latent feature values. The dendrogram was computed using Manhattan distances and Ward2 clustering and the latent features (columns) were converted to z-scores to depict relative differences between drug profiles. (C) Quantification of ATP content in drug-treated Mtb H37Rv cultures measured by the BacTiter-Glo™ luminescent enzymatic assay. Mtb was treated with DMSO, INH or INH in combination with four different doses at 0.1x-, 0.75x-, 5x- and 10x-IC50 of a second drug for 24 hours. The INH treatment dose was held constant at 10x IC50. Luminescence relative to culture OD600nm (RLU) is plotted (One-way ANOVA with Dunnett’s post-hoc test against INH+DMSO control: ***p < 0.001, **p < 0.01, *p < 0.05; ns, p > 0.05; n = 3). (D) Recovered CFU of 24-hour drug-treated Mtb H37Rv. Mtb was treated with DMSO, INH or INH in combination with the three doses used in (C), equivalent to 0.75x-, 5x- and 10x-IC50. After drug removal, cells were recovered on 7H10 agar plates and colonies were enumerated. The transparent point indicates that the CFU colonies were too dense to count for the data point. The recovery ratio is the Rx log_10_(CFU/mL) / DMSO control log_10_(CFU/mL), (Barplots depict the mean value and error bars are calculated from the standard error of the mean. One-way ANOVA with Dunnett’s post-hoc test against DMSO control: ***p < 0.001, **p < 0.01, *p < 0.05; ns, p > 0.05; n = 3).

Notably, fatty- and mycolic-acid biosynthesis gene expression were predicted to be most differentially expressed, suggestive of cell wall damage (Supplementary Figure 8C & 8D). Thus, an analysis of all the morphological and predicted transcriptional features highlights the cell wall effect of SQ109 (Supplementary Figure 8A). Yet, analyzing the same data using a feature-reduced space that is optimized for correlation with our RNA-seq dataset identifies a similarity to respiration inhibitors (Figure 5B). Together, this analysis suggests that the DECIPHAER latent space can reveal alternative insights into phenotypes given a single modality. These data also demonstrate that while the cell wall effect of SQ109 is dominant in each modality, SQ109’s profile in the DECIPHAER latent space is most similar to ATP-synthase inhibitors.

Next, we sought to identify the discriminating morphological and predicted transcriptional features contributing to SQ109-, bedaquiline-, and TBAJ-876-treated cells’ similarity in the DECIPHAER latent space. Across all morphological features, DNA staining intensity and variance in DNA-stain intensity were most significantly enriched, and elevated, in Mtb treated with the ATP synthase inhibitors and SQ109 (Supplementary Figure 9A, bottom). Though these nucleoid-related features may result from numerous physiological changes, previous studies in Mtb have demonstrated that nucleoid structure is linked to redox state^44^. Because bedaquiline, TBAJ-876, and SQ109 can interfere with respiration, an altered redox status might explain the increased DNA fluorescence in cells treated with these drugs. In support of this effect on respiration and redox status, bedaquiline, TBAJ-876, and SQ109 treatments were predicted to cause upregulation of folate biosynthesis, and nicotinamide and nicotinate metabolism and downregulation of the TCA cycle, consistent with a shift toward alternative energy production (Supplementary Figure 9B). Additionally, the Zur regulon, which is induced along with an altered respiratory and redox state^45^, was predicted to be the most significantly enriched pathway (Supplementary Figure 9B, top). Together, these data suggest that the similarity between bedaquiline, TBAJ-876, and SQ109 treatment responses in the DECIPHAER latent space is driven by their effects on respiration.

### SQ109’s respiration effect influences bacterial killing

Our model suggested that SQ109’s latent space signature was most similar to bedaquiline and TBAJ-876 due to an effect on respiration. Some MmpL3 inhibitors, like SQ109, have been shown to influence the proton motive force and respiration, but it is unknown if they also reduce ATP levels in Mtb^46^. Conversely, drugs that target the cell wall in Mtb, often increase cellular respiration and ATP levels^47,48^. To examine whether ATP depletion was a common phenotype among SQ109-, TBAJ-876-, and bedaquiline-treated Mtb, we quantified ATP within Mtb cultures in a previously established drug combination assay^47,48^. In this assay, the cell wall-acting wall acting agent, isoniazid, is used to induce an ATP burst, which can be diminished by adding a second antibiotic that inhibits ATP synthesis. As a control, we treated Mtb with bedaquiline which caused a dose dependent reduction in the ATP burst caused by isoniazid (Figure 5C). Adding the cell wall targeting antibiotic ethambutol to isoniazid further increased the ATP burst. Low doses of SQ109 (<5x IC50) resulted in increases in the ATP burst, like the effect of ethambutol. However, at high doses (> 5x IC50), SQ109 depleted ATP to levels similar to bedaquiline (Figure 5C).

To evaluate if ATP depletion in Mtb is a common phenotype among drugs that target MmpL3, we profiled two other MmpL3 inhibitors: BM212 and AU1235. Like SQ109, BM212 has been shown to dissipate the PMF in Mtb, whereas AU1235 does not^46^. Accordingly, BM212 caused a significant drop in the isoniazid-induced ATP burst at specifically high doses (> 5x IC50) (Figure 5C). In contrast, AU1235 caused a further increase to the ATP burst at high doses (Figure 5C). This suggests that the ATP depletion phenotype is shared only among some MmpL3 inhibitors and may be dependent on their ability to dissipate the PMF.

The phenotype of AU1235 suggested that inhibition of MmpL3 alone was insufficient to cause ATP depletion and that BM212’s and SQ109’s ATP depletion effect may thus be independent of their inhibition of MmpL3. To test this, we performed the ATP assay in an Mtb strain containing an anhydrotetracycline-inducible (ATc) MmpL3-F644L allele (Methods). This point mutation has previously been associated with partial SQ109 and AU1235 resistance^49^. We hypothesized that if the ATP depletion caused by the BM212 and SQ109 is independent of their inhibition of MmpL3, then the MmpL3-F644L inducible resistance mutant should still exhibit ATP depletion. Using the same drug concentrations as in the wildtype assay, SQ109 and BM212 still caused the isoniazid-induced ATP-burst to drop, suggesting that the ATP depletion is independent of MmpL3 inhibition (Supplementary Figure 10, Methods).

The isoniazid-induced ATP burst is thought to be lethal^47,48^. As such, adding increasing concentrations of a respiration inhibitor can rescue Mtb from isoniazid-induced cell death. We aimed to determine if the respiration effect of SQ109 and BM212 was dominant enough to rescue Mtb from isoniazid-induced cell death. The addition of the ATP-synthase inhibitor, bedaquiline, to isoniazid rescued Mtb from isoniazid-induced cell death, whereas adding the cell wall-acting drug, ethambutol, did not affect cell death (Figure 5D). We observed a dose-dependent increase in recovery of Mtb upon addition of SQ109 to isoniazid, similar to the greatest effect achieved by bedaquiline (Figure 5D). The addition of BM212 also recovered Mtb from isoniazid-induced cell death, albeit to a lesser degree than SQ109. These data suggest the inhibition of respiration by some MmpL3 inhibitors such as SQ109 and BM212 can influence Mtb cell death.

## DISCUSSION

Our goal in this study was to explore the mechanisms of drug action using the most important information from two modality measurements of Mtb drug response. To do this, we trained multi-modal autoencoders to compress transcriptomic and morphological Mtb drug response data into a shared latent space. Analysis of this space identified significant multi-modal features of cell damage, notably translation, DNA replication and repair, and cell size. We also discovered a novel correlated signature of damage involving phosphosugar stress and cell staining. Linezolid treatment disrupts homeostasis of this phosphosugar pathway which may contribute to the potency of the drug against Mtb. Our model generalized well to morphological data not included in the training phase, allowing us to make latent space- and transcriptional predictions from compounds for which we only measured one data modality. Using this function, which integrated data from low and high drug concentrations, we identified that Mtb treated with the MmpL3-inhibitor, SQ109, had a phenotype most similar to ATP-synthase inhibitors. We confirmed that high concentrations of SQ109 can deplete ATP levels in Mtb and that this effect on respiration can influence cell death. Together, our work establishes a quantitative framework, DECIPHAER, for determining root and proximal mechanisms of action by TB drugs.

Recent advancements in multi-omics integration technology have made it possible to map distinct modality measurements to a common latent space by training modality-specific autoencoders^23,24^. Such models have proven more powerful than previous methods as they can learn complex, nonlinear associations between modalities of dissimilar structure (images and RNA-seq) and, importantly in our study, do not require paired-modality measurements. Here, we included additional pre-training steps to these autoencoder-integration frameworks (Figure 1A) and extensively validated model inference (Figure 2 & Figure 4) which enabled us to explore and make predictions from drug data for which we had only a single modality measurement. Obtaining only one modality for certain biological samples is common in multi-omics datasets^13,50–53^ and our work demonstrates the utility of multi-modal autoencoders in these scenarios.

We utilized our model to identify correlated multi-modal features of damage. Many of these associations were consistent with the findings from a previous morphological study on CRISPRi-perturbed *M. smegmatis*^30^. Perhaps unexpectedly, we did not detect many strong correlations between cell size morphological features and expression levels of genes involved in cell elongation and division (e.g., Wag31 & LamA)^54^. Network analysis has suggested that antibiotics disrupt the coordination of transcriptional programs^55^, perhaps explaining the absence of correlations between Mtb cell size features and these genes’ expression levels. We hypothesize that if our dataset consisted of Mtb’s response to nutrient stress, to which it has evolved to overcome, these expected correlations might appear. The strong correlation between sugar metabolism and cell staining represents a novel Mtb cell damage signature (Figure 3A & 3C). Drugs that hinder protein synthesis, especially linezolid, seemed to influence the GlgE/Tres metabolic pathway and cell staining to the greatest effect (Figure 3C). Previous genomic screens have uncovered a protein synthesis and sugar metabolism co-vulnerability, providing further evidence that inhibiting protein synthesis may have downstream effects on sugar metabolism that lead to cell death^56,57^. Future analyses, perhaps in Mtb mutants, are required to conclusively determine whether sugar-phosphate toxicity contributes to protein synthesis inhibitors’ mechanism of Mtb cell death.

Predicting one modality from another is of great interest to scientists seeking to reduce experimental cost and time burden in their studies. DECIPHAER was able to accurately predict transcriptional signatures from imaging data alone (Figure 2 & Figure 4), suggesting that, after training, measuring only one modality may be required for multi-modal analysis of new TB drug profiles. The predicted transcriptional profiles of compounds for which we only measured morphological data were most similar to those of drugs that target the same broad cellular processes 65% of the time. This accuracy result was from an unsupervised learning task in which classification was not the primary goal and could be theoretically improved by implementing supervised learning. Even without supervision, we expect improved accuracy by including more drugs of diverse mechanisms in the training steps. Moreover, we hypothesize that the cases where predicted transcriptional data was not most like drugs targeting the same cellular process are due to the test drug having underexplored secondary mechanisms. For example, the predicted gene expression profile for the fatty acid-targeting drug, cerulenin, had the highest Jaccard similarity to DNA-acting agents (Supplementary Table 6). Cerulenin is known to inhibit topoisomerase activity in human cells^58^, and this same mechanism may contribute to the drug’s potency against Mtb. Surprisingly, the predicted transcriptional data for the beta-lactam antibiotic, D-cycloserine, was highly similar to the ETC-targeting drug clofazimine (Supplementary Table 6). A common mechanism of action that might explain the similarity in response of D-cycloserine and respiration drugs has not been reported and thus may be an impactful future direction.

Though SQ109 and BM212 have been shown to inhibit the PMF^46^, it has not been previously reported that they reduce ATP levels in Mtb. Given that these drugs also target the cell wall which can cause ATP bursts^47,48^, it was somewhat surprising that they reduced ATP levels in combinations with isoniazid and speaks to their respiration effects playing a dominant role in their mechanism. The effect of BM212 and SQ109 on respiration was potent enough to impact the viability of INH-treated Mtb (Figure 5D). This, along with the fact that SQ109 resistant mutants with mutations that map to MmpL3 are difficult to isolate^7^, suggests that SQ109’s effect on respiration influences Mtb killing and complicates the isolation of resistant mutants at high drug concentration. Moreover, our multi-modal latent space pointed to SQ109’s phenotype being most similar to respiration inhibitors, not cell wall-acting agents as expected based on conventional dimensionality reduction on single-modal data (Supplementary Figure 8A). This suggests that 1) inhibition of cellular respiration can be a phenotype-driving activity of some MmpL3 inhibitors and 2) DECIPHAER can highlight drug targets influencing killing of Mtb by identifying the important shared information from two modalities. The ability of the multi-modal space to highlight cell-death-relevant phenotypes is consistent with the burgeoning conclusion that multi-omics integration can improve insight by offering a more holistic view of a biological system than single-modal analysis^21,25,59,60^.

Several alterations to our study might improve DECIPHAER performance as well as our insights derived from the technology. Model architecture adjustments such as adding more hidden layers or nodes to the autoencoder architecture might improve the integration and cross-modal predictions. We opted for more shallow neural networks in this study because optimizing many parameters through training on a relatively small dataset size (by *n*) can result in overfitting^61^. Thus, using a deeper architecture would likely require more drug treatments, growth conditions, or treatment time-points to sample more of the morphological and RNA-seq spaces and improve insights. Adding other modality measurements, such as gene-level fitness measurements (CRISPR screens), metabolomics, or additional fluorescent cell stains, could help link drug-induced phenotypes to functional consequences. Such datasets could even be integrated as a third modality, as there is theoretically no limit to the number of modality-specific autoencoders that can be trained for integration with our multi-modal latent space. As training data sizes increase, we anticipate the DECIPHAER platform will enable researchers to determine the most critical mechanisms of novel TB drugs and make cross-modal predictions from only single-modal measurements.

## METHODOLOGY

### Bacterial strains and culture conditions

For the MmpL3 ATP and CFU assays, *M. tuberculosis* H37Rv transformed with the plasmid pGMCK-T10M-P606-mmpL3-F644L-TsE5-bc was used. This plasmid contains the MmpL3-F644L allele which is overexpressed upon ATc exposure. For RNA-seq and RNA-seq IC determinations, *M. tuberculosis* Erdman containing plasmid pMV306hsp+LuxG13 (PMID: 34469743) was used, as these experiments were originally designed to match other orthogonal studies performed in the strain background. For all other assays, wild type (WT) *M. tuberculosis* Erdman cells were used. In all experiments, cells were adapted to either standard growth or cholesterol media as indicated by figure legends. Mtb was stored at -80°C in standard medium in 1 mL aliquots at mid-log phase (OD_600_ ∼0.5-0.7). Prior to culture media adaptation, these frozen aliquots were subcultured to OD_600_ = 0.05 in standard media. Then, once these reached mid-log phase, the cultures were diluted in their respective media to begin adaptation. All media was buffered with 100 mM 3-(N-morpholino)propanesulfonic acid at pH 7. Standard growth medium consisted of 7H9 broth, (ThermoFisher; DF0713-17-9), 0.2% glycerol, 10% OADC (0.5g/L oleic acid, 50g/L albumin, 20g/L dextrose and 0.04g/L catalase), and 0.05% Tween-80. For maintenance of the MmpL3-F644L overexpression strain, the standard growth medium was supplemented with 25 μg/mL of the selection antibiotic, kanamycin, throughout all culture steps except during drug treatment. For cholesterol growth medium, a base medium consisting of 7H9 powder (4.7g/L), fatty acid-free BSA (0.5g/L), NaCl (100mM) and tyloxapol (0.05%) was made. Within 48 hours prior to cholesterol medium cell culturing, cholesterol was dissolved in a 1:1 (v/v) mixture of tyloxopol and ethanol at a concentration of 100mM and heated to 80°C for 30 minutes. This stock was then added to pre-warmed (37°C) base medium to a final cholesterol concentration of 0.05 mM^62^. All liquid cultures were grown at 37°C in vented 25 mL tissue-culture flasks with shaking at 100 RPM until transferring to 96-or 384-well plates for drug treatments.

### Drug treatments

The compounds used for this study are listed in Supplementary Table 1 along with their corresponding solvents, inhibitory concentrations and literature-reported targets. Drugs were dispensed with an HP D300e digital dispenser in 96 or 384-well plates. For drugs dissolved in water, Triton-X100 was added to a final concentration of 0.01% to reduce surface tension and promote accurate volume dispensing. The final DMSO concentration was kept below 1.5% for all drug treatments. For inhibitory concentration (IC) assays, the locations of all drugs in the wells were randomized to prevent any plate-location effects. Culture volumes for 96- and 384-well plates were 150 μL and 50 μL per well, respectively. The outer wells of all plates did not contain culture but were instead filled with an equal volume of media. All drug treatment plates were kept at 37°C in humidified plastic bags to reduce evaporation.

### Plate measurements

Plate measurements were performed in 384-well plates using the BioTek Synergy Neo2 Hybrid Multi-Mode Reader. For IC determination assays, measurement time-points were chosen to be ∼5-8 doubling times after drug exposure. This meant that for experiments in the standard growth condition, plates were read on day 7 whereas for experiments in the cholesterol growth condition, plates were read after 14 days. Drugs that did not reach an IC50 value were considered inactive. Plates from the ATP quantification assays were read after 22 hours of drug exposure.

### RNA-seq library preparation and data pre-processing

Mtb strain Erdman with plasmid pMV306hsp+LuxG13 was adapted to the acidic, cholesterol and valerate growth conditions as previously described^62^ prior to drug exposure and RNA extraction. Briefly, Mtb was first cultured in the standard growth medium and then subcultured at least once into the designated growth conditions prior to drug treatment. Once the bacterial cultures reached an OD_600nm_ of ∼0.5, the cultures were diluted to an OD_600nm_ of 0.05 in 50ml conical tubes containing drug at a concentration of 1x the IC90. Drug-treated cultures and untreated controls were then harvested by centrifugation at time points 4- and 24-hours post-drug treatment. To halt transcriptional responses during the RNA extraction process, GTC buffer was added to the cultures, and they were washed twice in GTC buffer. The resulting pellets were stored in Eppendorf tubes containing ∼1ml of GTC buffer (4M guanidinium thiocyanate) + 0.5% sarcosyl, 25 mM sodium citrate and 0.1M beta-mercaptoethanol. Samples were stored in aerosol-tight O-ring caps at -80°C until RNA extractions were performed. On the day of extraction, the frozen GTC-treated pellets were thawed at 37 degrees, harvested by centrifugation, resuspended in prewarmed triazole and transferred to 2ml O-ring tubes containing sterile 0.2 mm zirconia beads. The triazole mixture was then bead beaten twice followed by the addition of chloroform and vigorous shaking for one minute. The samples were then centrifuged, and the aqueous layer was removed and placed in Qiagen RNeasy columns for RNA cleanup. Genomic DNA was removed from all samples by performing a DNase digest on the columns before elution in pre-warmed RNase free water. The extracted RNA was then removed from the BSL-3 facility. Concentration and quality of RNA (260nm/280nm absorbance) was assessed using a nanodrop spectrophotometer and stored at -80°C until sample submission. Samples were submitted to the Microbial ‘Omics Core (MOCP) and Genomics Platform at the Broad Institute of MIT and Harvard. QC, library preparation, ribodepletion, and paired-end sequencing on the Illumina NovaSeq 6000 following a modified RNAtag-Seq method^63^ was performed by the MOCP.

Sequencing reads were aligned to the *Mycobacterium tuberculosis* str. Erdman = ATCC 35801 (RefSeq assembly accession: GCF_000350205.1) using BWA^64^. Reads were then sorted and indexed with samtools (v1.9) using the commands “samtools sort -l 0” and “samtools index”. Gene-level counts were obtained from the sorted, genome-mapped reads using the python-based package HTSeq (v0.13.5). Genes with counts less than an average of one were removed and samples with fewer than 500,000 reads were dropped. To adjust for unwanted variation stemming from the two sequencing batches, the count matrix was adjusted using ComBat-seq batch correction (v1.32.0). The read count matrix was then normalized using DESeq2’s (v1.32.0) variance stabilizing transformation (VST) to account for sequencing depth and ensure homoscedasticity across the transcriptome. Erdman locus tags were converted to their H37Rv equivalent using a custom annotation table which was derived from the mycobrowser database and generously provided by Shumin Tan (Tufts University). The resulting gene expression matrix was used for autoencoder training.

### Morphological profiling assay

Mtb Erdman cells were adapted to the standard growth media for 3-5 doubling times. They were subcultured once more for 5-6 doubling times, until reaching an OD_600_ of 1.2. Subsequently, the cultures were diluted to an OD_600_ of 0.3 in 96-well plates containing pre-printed drugs. Drug concentrations were 0.75x and 10x the IC50 if they were single treatment or, if it was a combination treatment, 1x the IC50 value of each drug treatment. After 17 hours of drug treatment, Mtb cells were fixed in the 96-well plates using a final concentration of 4% paraformaldehyde (ThermoFisher; 043368-9M). Cells were pelleted and washed twice with 100 μL of PBS (ThermoFisher; 20012-027) + 0.2% Tween-80 (PBST), then resuspended with 100 μL of PBST for storage at 4 °C. Following fixation, staining was performed with 0.6 μg of FM4-64FX (ThermoFisher; F34653) and 15 μL of a 0.1 μM SYTO 24 (ThermoFisher; S7559) stock in each well containing PBST and fixed bacilli. The plate was then incubated at room temperature in the dark for 30 min. Once stained, the cells were washed once with an equal volume of PBST and resuspended in 30 μL of PBST. Stained Mtb was spotted onto agar pads (2% wt/vol agarose; SigmaAldrich; A3643-25G). Images were captured with a widefield DeltaVision PersonalDV (Applied Precisions) microscope and DV Elite CMOS camera. Bacteria were illuminated using an InsightSSI Solid State Illumination system with transmitted light for phase contrast microscopy. SYTO 24 signal was captured using Ex. 475 nm and Em. 525 nm, and FM4-64FX with Ex. 475 nm and Em. 679 nm. Montage images were generated using a custom macro that captures 25 individual fields of view per image.

### Single-cell segmentation and feature calculations

To generate single-cell profiles of morphological features, microscopy images were processed with a custom analysis pipeline. First, we performed an illumination correction using BaSiC^65^ to ensure all channels have an even distribution of illumination across the dimensions of each image. To segment single-cell boundaries in images and extract morphological measurements we utilized the ImageJ plugin, MicrobeJ (v5.13I). The pipeline generated a matrix of 25 morphological measurements across all cells (∼1.14 million cells). Because of the subtle and heterogeneous morphological response of Mtb to drug treatment, clustering of single cells by their morphological response does not discriminate disparate drug treatments. To address this, we used summary statistics of features from subsets of cells to generate robust morphological profiles. For each perturbation (single-drug or drug-combination), 1000 cells at a time were randomly sampled without replacement and the mean, first quartile, and third quartile were calculated for each morphological feature. Sampling was repeated until the number of unused single cells was exhausted. After excluding features with a standard deviation of 0, we obtained 78 summary statistic features from these bulk measurements. The summary statistic features were used as input for drug-phenotype clustering and autoencoder training. Using cell summaries instead of single-cell data increased the F-test statistic, or the ratio of between-group variance to within-group variance, of each drug treatment group. The high F-test statistic indicates the morphological profiling data points cluster strongly according to their drug treatment.

### DECIPHAER framework

Here, we formalize our approach for training multiple variational-autoencoders on two modality measurements of Mtb drug response. Let *n* represent the number of features in each modality’s matrix, *X* and *Y*. Many of these are “noise features”, or features that do not discriminate phenotypes of interest in the latent space and have no correlation with the other modality’s set of features.

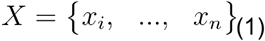

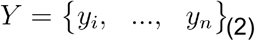

The encoder modules of each autoencoder *f*_*e*_(*X*) and *g*_*e*_(*Y*) determine a latent representation *z* which can be mapped from *X* and *Y*.

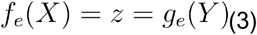

The decoder modules of each autoencoder *f*_*d*_(*z*) and *g*_*d*_(*z*) take in the same latent representation and estimate each modality’s matrix, *X′* and *Y′*, respectively.

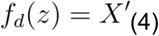

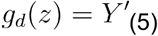

*θ*_*i*_ and *γ*_*i*_ are the learnable coefficients (weights/biases of the modality specific autoencoders) for each i feature in each respective data modality. These coefficients effectively determine the importance of each feature for encoding z. Thus, an optimal latent space should correspond to low weights for *θ*_*i*_ or *γ*_*i*_ if i is a “noise feature”.

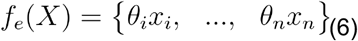

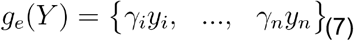

To optimize the *θ*_*i*_ and *γ*_*i*_ coefficients of the autoencoders, we employ the following loss functions proposed by Dai Yang & Belyaeva et al^23^. For modality X’s autoencoder, we optimize through gradient descent:

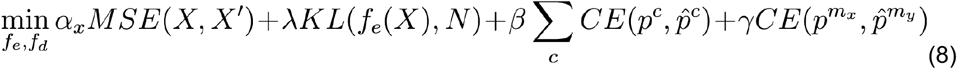

For modality *Y*’s autoencoder, we optimize the similar objective function through gradient descent:

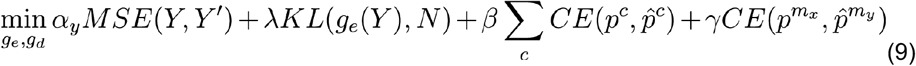

where the *α, λ, β* and *γ* coefficients correspond to the user-defined hyperparameter weights for each of the terms in the loss function. All terms are summed and the autoencoders are trained to minimize this sum. The *α* and *λ* hyperparameters are the weights for typical variational autoencoder loss. They correspond to the reconstruction (quantified by mean-squared error or MSE) and Kullback-Leibler (KL) divergence losses, respectively. The reconstruction loss ensures the latent space holds information that describes the features, and relationships between features, in each data modality. The KL divergence loss quantifies the difference between the encoding distribution *f*_*e*_(*X*) and the standard normal distribution *N*. High values for the *α* term ensure accurate reconstruction from the latent space whereas high values for the lambda term promote regularization.

To ensure the latent space contains information that discriminates disparate drug phenotypes and clusters similar ones, we employ a drug phenotype classifier loss term, controlled by the *β* hyperparameter above. We use categorical drug phenotype labels *c* to train a linear classifier with the terms *A* and *b* to predict the drug phenotype 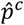, given the latent embedding *z* of *X* or *Y*’s data points.

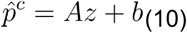

The autoencoder’s objective functions (8 & 9) minimizes the multi-class cross-entropy (CE) of the predicted drug phenotype 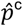 and the true drug *p*^c^ across the latent embeddings for modality *X* and *Y*. Thus, high weights for the beta hyperparameter ensures data points of similar drug phenotypes are clustered together in the latent space, regardless of the data modality from which they were sampled.

To further ensure the latent embedding contains no modality-specific information, we utilize the discriminative loss term, controlled by the *γ* hyperparameter. We train a classifier *f*(*z*) to predict the modality 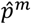 of each data point given its latent embedding.

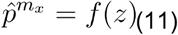

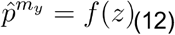

*f*(*z*) is optimized by minimizing the binary CE between the predicted and true modality label.

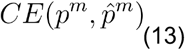

The discriminative loss terms in 8 and 9, used to train the autoencoders, quantifies the CE between the modality probabilities predicted for *X*, 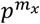, and its true modality labels of *Y*,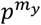. In other words, the autoencoders are trained to fool the modality discriminator. Thus, high values for *γ* ensure the latent space contains minimal modality specific information.

### DECIPHAER training for the integration of RNA-seq and morphological data

For DECIPHAER integration training, we subsetted the modality datasets to include only the drug treatments for which we had both RNA-seq and morphological data (Supplementary Figure 1). This resulted in 15 treatment groups, including untreated, consisting of RNA-seq from two time-points per drug in the cholesterol growth condition and morphological data from one time-point in the standard growth condition. All gene expression- and morphological features were subtracted by their mean and divided by their standard deviation (z-scored) prior to training. For model evaluation, each dataset’s held-out test splits were divided using stratified 6-fold cross-validation (sklearn, v0.0.post1) and the scalar calculated from the corresponding training split was used to z-score the data. Unless noted otherwise in the main text, the hyperparameters used to train DECIPHAER are listed in Supplementary Table 1 and were chosen empirically based on test performance through full-factorial sampling. The DECIPHAER network architecture we used throughout this work is displayed in Supplementary Table 2.

### Clustering to define drug-phenotype labels

A key component of the DECIPHAER integration tool is the drug-phenotype latent space classifier which relies on labeled data. This classifier takes a data point’s latent space coordinates as input and predicts its drug-phenotype label as output (Equation 6) and ensures that similar drug treatment measurements have small distances in the latent space. Before training the cross-modal autoencoders on our morphological and transcriptional data, we sought to generate accurate drug-phenotype labels. The morphological profiling data was chosen over the RNA-seq dataset for defining drug-phenotype labels because of its lower dimensionality and higher “n”, both of which result in better clustering performance. We developed a two-prong aggregation method for defining cluster labels to balance the tradeoffs between clustering algorithms. Specifically, we utilized two algorithms, HDBScan (dbscan v0.0.12, parameters) on UMAP embeddings, and Leiden community detection (LCD) (scanpy v1.9.3) on PCA embeddings from the first 5 PCs to define clusters. All data points from each drug treatment were assigned to the drug’s most popular cluster label from each clustering algorithm. To aggregate the HDBScan and LCD results, drug treatments that were assigned to the same cluster in one algorithm but different clusters in the other were given new labels. This method revealed 11 clusters (distinct drug-response phenotypes), the labels of which were used to train the drug-phenotype latent space classifier on the two modalities.

### Autoencoder pretraining

To ensure DECIPHAER learned patterns of morphological and gene expression features that would generalize, the RNA-seq and morphological autoencoders were pre-trained with separate latent spaces before training the two networks on a shared latent space. The data used for pretraining consisted of drug treatment response data for which only one modality measurement was collected (Supplementary Figure 1 and Supplementary Table 1) and was z-scored using the mean and standard deviation of the training set. For the morphological pre-training-set, this consisted of 38 different single and two-way-combination drug treatments. The RNA-seq autoencoder pre-training-set included the same 14 drugs used for DECIPHAER integration, but in the valerate and acidic growth conditions. It also contained RNA-seq responses from three additional compounds from the acidic, valerate and cholesterol growth conditions for which we did not have matched morphological data (Supplementary Figure 1). The morphological and RNA-seq autoencoders were pretrained on 50-dimensional latent spaces for 100 and 400 epochs, respectively, using the KL divergence and reconstruction losses.

### RNA-seq geneset calculations

To define sets of differentially expressed genes in each drug treatment’s true RNA-seq data, we used the function “deseq” in R (DESeq2, v1.40.2). For each drug treatment, we grouped the 4- and 24-hour time points together and calculated adjusted p-values and fold changes relative to untreated of the same time points. We selected marker genes for each drug by first selecting the 300 genes with lowest adjusted p-values then subsetting from this list the 100 genes with greatest fold-changes. These lists were used to evaluate model predictions as in Figure 2.

### Jaccard similarity coefficient for predicted transcriptional profiles

To compare the transcriptional signatures predicted from morphological profiles of drugs for which we did not measure RNA-seq to our training set, we utilized the Jaccard similarity coefficient which quantifies similarity between two qualitative sets. The Jaccard similarity coefficient between genesets *a* and *b* is defined as the number of elements in the intersection divided by the number of elements in the union:

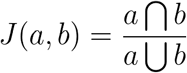

Specifically, we first generated robust predicted transcriptional profiles by translating all morphological data 920 times and selecting the median gene expression values of each data point. These iterations minimize aberrant predictions stemming from the random nature of the variational autoencoders. We could not use DESeq2 to calculate differentially expressed genes for predicted data as it had been transformed significantly for DECIPHAER input and violated the assumptions of the “deseq” function. Instead, we used Wilcoxon rank-sum tests comparing each drug treatments’ predicted differential expression against the predicted differential expression profile of all training-set drug treatments and untreated. Finally, using these differential expression sets, we generated an average similarity metric between each drug’s differential expression list and the training-set drugs’ lists. To calculate this average, we took the mean of the Jaccard similarity coefficient of every possible equal-sized set in the 50 to 500 genes of lowest p-value (450 set comparisons for each pair of drugs). Through this set comparison approach, we avoid some of the issues that arise when attempting to calculate drug-drug transcriptional distances in the high-dimensional RNA-seq space.

### Gene expression pathway enrichments

To quantify pathway enrichment, we utilized the PathFindR software in R (pathfindr, v2.3.1) which identifies enriched subnetworks of genes based upon their expression level and a protein-protein interaction network. We utilized a custom gene:pathway annotation file derived from Kyoto Encyclopedia of Genes and Genomes (KEGG) and literature searches.

### Metabolomics

Targeted metabolomics experiments were performed using previously described methods^66,67^. In brief, 1 ml of Mtb Erdman culture from the mid-log phase was passed through 0.22 μm nylon filters and cultured for 5 days at 37 °C on 7H10 agar media (BD) supplemented with dextrose and glycerol. On day 6, these bacteria-laden filters were transferred onto reservoirs filled with cholesterol (200 μM) containing media (no tyloxapol) and allowed to grow for the next 3 days to acclimatize to cholesterol as a new carbon source. These cholesterol-adapted bacteria were transferred onto new reservoirs containing 1x, 5x, and 10x IC50 doses of different antibiotics for 48 hrs and then collected in 1 ml precooled mixture of acetonitrile: methanol: water (40:40:20). Cells were lysed using 0.1 mm Zirconia beads and extracted metabolites were further processed to profile on LC-MS. Two methods of liquid chromatography were used (i) formic acid and (ii) hydrophilic interaction liquid chromatography (HILIC). For the formic acid method, 2 μl of every samples were resolved on a Diamond Hydride Type C Column (Cogent) using 1260 Infinity liquid chromatography (Agilent) coupled to an Agilent Accurate-Mass 6230 TOF-MS (both positive and negative modes). For the liquid phase, solvent A (water with 0.2% formic acid) and solvent B (acetonitrile with 0.2% formic acid) were used at the following gradients with a flow rate of 0.4 ml min−1: 85% B (0–2 min); 80% B (3–5 min); 75% B (6–7 min); 70% B (8– 9 min); 50% B (10–11 min); 20% B (11–14 min); 5% B (14–24 min) and 10 min of re-equilibration period using 85% B. While for HILIC 3 μl of the extracted lysates were separated on Agilent 1290 Infinity liquid chromatography through an Agilent InfinityLab Poroshell 120 HILIC-Z, 2.1 × 150 mm, 2.7 μm column heated to 50 °C. Solvent A (100% H_2_O with 10 mm ammonium acetate, 5 mM InfinityLab Deactivator Additive, pH 9 using NH_4_OH) and Solvent B (85% acetonitrile in H_2_O with 10plotted mM ammonium acetate, 5 mM InfinityLab Deactivator Additive, pH 9 using NH_4_OH) were used at a flow rate of 0.250 ml/min with the following gradient: 96% B for 0–2 min; 88% B for 5.5–8.5 min; 86% B for 9–14 min; 82% B for 17 min, 65% B for 23–24 min; 96% B for 24.5–26 min; and the end-run at 96% B for 10 min. Results were collected on an Agilent 6230 TOF mass spectrometer operated at 3500 V Cap and 0 V nozzle voltage in extended dynamic range, negative mode. All results were collected in centroid mode for m/z values from 50 to 1700. Ion abundances were estimated using Profinder 8.0 (Agilent Technologies). Standard metabolites were used to confirm the accuracy of identified peaks. Absolute and relative abundances (with respect to the untreated control) were plotted.

### ATc-inducible MmpL3-F644L mutant construction

Plasmid pGMCK-T10M-P606-mmpL3-F644L-TsE5-bc, which allows ATc-inducible expression of the MmpL3-F644L allele was obtained from GenScript. In this plasmid, transcription of MmpL3-F644L is mediated by the promoter by a strong promoter (P606), which carries a single tet operator (tetO) between its -10 and -35 elements and is repressed a Tet repressor, TetR10, also encoded within the plasmid. The plasmid was transformed into Mtb H37Rv with selection on Middlebrook 7H10 (BD Biosciences) agar plates containing 10% OADC (BD Biosciences), 0.5% glycerol (Sigma) and 25 μg/ml kanamycin (Sigma). The plates were incubated at 37°C for 21 days. Transformants were inoculated and cultured in Middlebrook 7H9 (BD Biosciences) containing 10% ADNaCl, 0.2% glycerol, 0.05% Tyloxapol (Sigma) and 25 μg/ml kanamycin. The presence of the MmpL3-F644L allele was confirmed by PCR amplification and Sanger sequencing. To confirm the strain exhibits an increased resistance to SQ109, an inhibitory concentration (IC) assay was performed. SQ109 was dissolved in DMSO and dispensed with an HP D300e digital dispenser into 384-well plates. All strains were cultured in 7H9 containing 10% ADNaCl, 0.2% glycerol, 0.05% Tyloxapol and, as needed, 25 μg/ml kanamycin, both with and without 500 ng/mL of ATc. Once the bacterial cultures reached an OD580nm of ∼0.6-0.8, the cultures were diluted to an OD580nm of 0.01 in 15ml conical tubes. The cultures were added to the drug containing 384-well plates and incubated at 37°C. The plates were read after 14 days on a Molecular Devices SpectraMax M2 microplate reader. Analysis of the results indicated that the H37Rv::MmpL3-F644L strain exhibited an 8-fold increase in the IC50 of SQ109 in the presence of ATc compared to wild-type H37Rv.

### ATP assay

WT Mtb H37Rv or Mtb MmpL3-F644L-overexpression cells were adapted to the standard growth media for 3-4 doubling times. Each strain was subcultured once more at which time the MmpL3-F644L allele was induced for overexpression with 500 ng/mL of ATc. The cultures were grown for 4.5 doublings, at which point the OD_600_ = 0.6-0.7 cultures were diluted to OD_600_ = 0.1 in 384-well plates containing pre-printed drugs. After 22 hours of drug treatment, OD_600_ was measured for each plate. This time point was designed to be short so as to avoid the effects of cell death and growth inhibition on the ATP read-out^48^. To quantify ATP levels, we utilized a commercially available luminescent assay which relies on a bacterial cell lysis step followed by a recombinant luciferase enzymatic reaction. Briefly, we added 50 μL of BacTiter-Glo™ (Promega; G8230) to an equal volume of drug-treated cell culture in each well. Plates were rocked gently by hand for two minutes then incubated at 37°C for five minutes. Finally, the plates were rocked gently by hand for another three minutes and their luminescent signals, which are proportional to the amount of ATP present, were measured. Each well’s luminescent signal was divided by its post-drug treatment OD_600_ value to obtain relative luminescent units (RLU). All ATP assays were performed in experimental triplicate.

### Mtb recovery assay

Colony forming unit (CFU) assays to quantify recovery after drug treatment were designed to match the protocol of the ATP assay as close as possible. WT Mtb H37Rv cells were adapted to the standard growth media for 3-4 doubling times and subsequently subcultured for 4.5 doubling times. The cultures, with a density of OD_600_ = 0.7, were diluted to OD_600_ = 0.1 in 96-well plates containing pre-printed drugs. After 22 hours of drug treatment, Mtb cells were washed twice with 120 μL PBST to remove drugs and pellets were resuspended to a volume of 150 μL PBST per well. The washed cells were serially diluted in PBST and the 10^−2^, 10^−3^, 10^−4^ and 10^−5^ dilutions were spread on quarter plates containing sterilized 7H10 agar (Millipore-Sigma; M0303-500G) + 10% OADC + 0.2% glycerol. Plates were incubated at 37°C for three weeks at which point colonies were counted. CFU assays were performed in experimental triplicate.

### Computational analyses

We used both python and R for computational analysis. For data reading and writing, we used the R packages readr (v2.1.5), readxl (v1.4.3) and the python package, pandas (v1.5.3). Data munging in R was performed using the packages tidyverse (v2.0.0), stringr (v1.5.1), dplyr (v1.1.4) and purrr (1.0.2). Data munging in python was performed using pandas (v1.5.3). UMAP and PCA were computed using the R functions umap (umap; v0.2.10.0) and prcomp (stats; v4.3.1). DECIPHAER modelling was performed in python mainly using pytorch (torch; v1.13.1), along with a few functions requiring sklearn (v0.0.post1) and numpy (v1.23.5). For kNN analysis, we utilized the R package class (v7.3-22) with Euclidean distance on UMAP embeddings. For hierarchical clustering and heatmap plotting, we used pheatmap (v1.0.12). For all other plotting we used the R packages ggplot2 (v3.4.4) and ggpubr (v0.6.0).

### Statistical analysis

All statistical analysis was performed in R and python. To compute enriched genes in predicted RNA-seq and enriched morphological features, we used the Wilcoxon rank-sum test in the python package scipy (v1.10.0). For the Wilcoxon rank-sum test in Supplementary Figure 5, we used the function wilcox_test from the R package rstatix (v0.7.2). ANOVA and Dunnett’s tests were performed in R with the stats package and DescTools (v0.99.54), respectively. T-tests were computed using rstatix. Pearson’s and Spearman’s rank correlation coefficients were computed using the stats package in R. We also used the stats package to compute the F-test with the following formula:

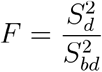

where *S*_*bd*_^2^ represents the variance between drug-phenotype clusters and the *S*_*d*_^2^ variance within drug-phenotype clusters.

## DATA AND CODE AVAILABILITY

A basic version of the code used for training and testing of DECIPHAER is available at the Github repository: wjohnsonTufts/deciphaer.

## ACKNOWLEDGEMENTS

We thank members of the Aldridge laboratory, R. Isberg, S. Tan, and L. Hu for detailed discussion. This work was supported in part by the Bill & Melinda Gates Foundation under grants INV-010515, INV-007519 and INV-027276 awarded to BBA, INV-004761 to DS, and INV-027106 to PKS. We thank GlaxoSmithKline (GSK) for providing GSK-839, GSK-286 and sanfetrinem, the TB Alliance for providing TBAJ-876, TBA-7371 and TBI-223, the University of Dundee for providing the lysyl-tRNA synthetase inhibitor, and Otsuka Pharmaceutical for providing quabodepistat.

## DECLARATION OF INTERESTS

The authors declare no competing interests.

## Notes

### Competing Interest Statement

The authors have declared no competing interest.

