## Supplementary Data for "Integration of multi-modal measurements identifies critical mechanisms of tuberculosis drug action"

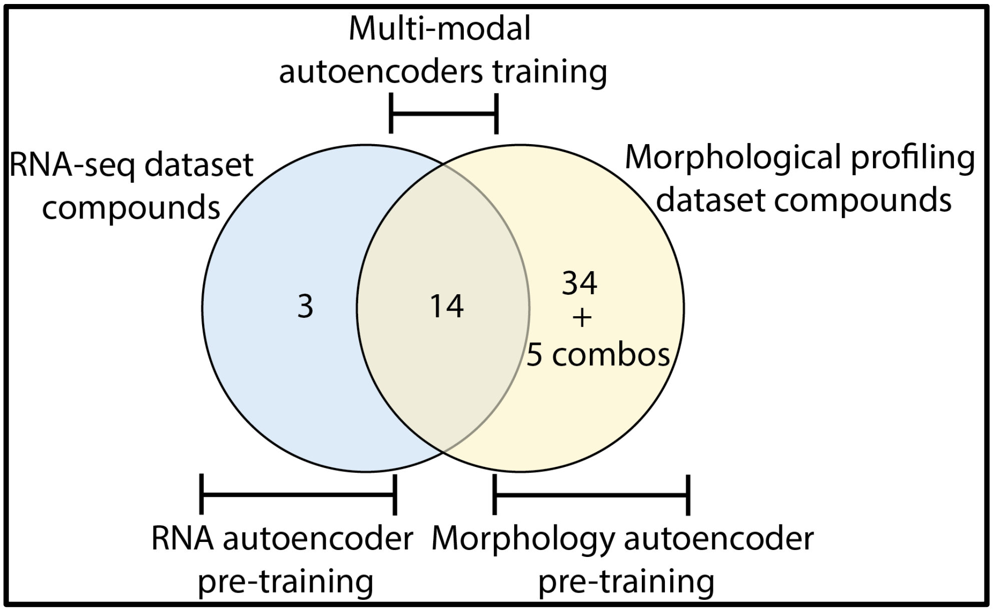


Supplementary Figure 1**. Quantities of drug treatments used to generate the datasets used for DECIPHAER training**. To generate an RNA-seq dataset, Mtb was adapted to three growth conditions and treated with 17 different drugs for two different treatment durations. The morphological profiling dataset consisted of 14 drug treatments that match the RNA-seq dataset along with 34 other drug treatments and 5 combination antibiotic treatments. The data from the 14 drug treatments present in each dataset are used for DECIPHAER alignment training whereas drug treatments not present in both datasets are used for DECIPHAER pretraining and model inference.

| **Drug Name** | **Drug Code** | **Solvent** | **Vendor/Source** | **Catalog number** | **RNA-seq drug vendor/source** | **RNA-seq drug catalog** | **Have RNA-seq data** | **Have morphological data** | **IC50 in standard for morphological profiling (µg/mL)** | **IC90 in cholesterol for RNA-seq (µg/mL)** | **Cellular Target** |
|---|---|---|---|---|---|---|---|---|---|---|---|
| bedaquiline | BDQ | DMSO | Asta Tech | 43211 | Sigma Aldrich | ATE517247923 | Yes | Yes | 1.47E-01 | 7.16E-02 | Respiration |
| clofazimine | CLZ | DMSO | SigmaAldrich | C8895 | Sigma Aldrich | C8895 | Yes | Yes | 6.15E+00 | 2.16E+00 | Respiration |
| ethambutol | EMB | DMSO | Alfa Aesar | J60695 | Sigma Aldrich | E4630 | Yes | Yes | 2.61E+00 | 7.88E+00 | Cell wall |
| isoniazid | INH | DMSO | SigmaAldrich | I3377 | Sigma Aldrich | I3377 | Yes | Yes | 2.63E-02 | 4.00E-02 | Cell wall |
| lysyl-tRNA synthetase inhibitor | KRS | DMSO | University of Dundee | Gift | University of Dundee | Gift | Yes | Yes | 2.24E-01 | 1.70E-01 | Protein |
| linezolid | LIN | DMSO | Apex Bio | A5181 | Sigma Aldrich | PZ0014 | Yes | Yes | 9.02E-01 | 1.72E+01 | Protein |
| moxifloxacin | MOX | DMSO | Alfa Aesar | J66626 | Sigma Aldrich | SML1581 | Yes | Yes | 2.15E-01 | 1.64E+00 | DNA |
| pretomanid | PRE | DMSO | Apex Bio | A1736 | Apex Bio | A1736 | Yes | Yes | 9.66E-02 | 1.66E-01 | Cell wall |
| pyrazinamide | PZA | DMSO | TCI Chemicals | P0633 | Sigma Aldrich | PHR1576 | Yes | Yes | Not reached | 1.18E+02 | Respiration |
| rifapentine | RFP | DMSO | Apex Bio | B2127 | Sigma Aldrich | R0533 | Yes | Yes | 1.31E-02 | 8.60E-02 | RNA |

| | | | | | | | | | | | |
|---|---|---|---|---|---|---|---|---|---|---|---|
| rifampicin | RIF | DMSO | TCI Chemicals | R0079 | Sigma Aldrich | R3501 | Yes | Yes | 4.02E-02 | 2.75E-01 | RNA |
| tbaj-876 | SFN | DMSO | TB Alliance | Gift | TB Alliance | Gift | Yes | Yes | 1.28E-02 | 1.00E-02 | Respiration |
| tba-7371 | TBA | DMSO | TB Alliance | Gift | TB Alliance | Gift | Yes | Yes | 1.55E+00 | 2.14E-01 | Cell wall |
| tbi-223 | TBI | DMSO | TB Alliance | Gift | TB Alliance | Gift | Yes | Yes | 3.04E+00 | 2.01E+00 | Protein |
| gsk-839 | GS8 | DMSO | NA | NA | GSK | Gift | Yes | No | NA | 2.00E-03 | Protein |
| gsk-286 | GS2 | DMSO | NA | NA | GSK | Gift | Yes | No | NA | 5.00E-03 | Unknown |
| sanfetrinem | SAN | Water | NA | NA | GSK | Gift | Yes | No | NA | 7.80E-01 | Cell wall |
| actinomycin D | ACT | DMSO | Fisher Scientific | AAJ60148LB0 | NA | NA | No | Yes | 2.79E+00 | NA | RNA |
| amikacin | AMK | Water | SigmaAldrich | A3650 | NA | NA | No | Yes | 1.50E+00 | NA | Protein |
| ampicillin | AMP | DMSO | SigmaAldrich | A9393 | NA | NA | No | Yes | 2.50E+01 | NA | Cell wall |
| chloramphenicol | CAM | DMSO | SigmaAldrich | C0378 | NA | NA | No | Yes | 4.82E+00 | NA | Protein |
| capreomycin | CAP | Water | Fisher Scientific | ICN15492483 | NA | NA | No | Yes | 4.39E-01 | NA | Protein |
| CCCP | CCC | DMSO | SigmaAldrich | C2759 | NA | NA | No | Yes | 1.90E+00 | NA | Respiration |
| cerulenin | CER | DMSO | SigmaAldrich | C2389-10mg | NA | NA | No | Yes | 2.93E+00 | NA | Cell wall |
| ciprofloxacin | CIP | Water | SigmaAldrich | phr1044-1g | NA | NA | No | Yes | 1.41E-01 | NA | DNA |
| clarithromycin | CLA | DMSO | SigmaAldrich | A3487-100MG | NA | NA | No | Yes | 1.07E+00 | NA | Protein |
| cefotaxime | CTA | Water | SigmaAldrich | C7039 | NA | NA | No | Yes | 9.10E+00 | NA | Cell wall |
| cycloserine | CYC | DMSO | Fisher Scientific | 239831 | NA | NA | No | Yes | 1.82E+00 | NA | Cell wall |
| daunorubicin | DAU | DMSO | SigmaAldrich | 30450-5MG | NA | NA | No | Yes | 3.07E+00 | NA | DNA |
| delamanid | DEL | DMSO | Advanced ChemBlocksInc | L13485 | NA | NA | No | Yes | 3.65E-03 | NA | Cell wall |
| doxycycline | DOX | DMSO | SigmaAldrich | D9891 | NA | NA | No | Yes | 3.44E+00 | NA | Protein |
| ethionamide | ETA | DMSO | TCI Chemicals | E0695 | NA | NA | No | Yes | 6.75E-01 | NA | Cell wall |
| fidaxomicin | FID | DMSO | Fisher Scientific | 50-101-5249 | NA | NA | No | Yes | 2.74E+00 | NA | RNA |
| gentamicin | GEN | Water | SigmaAldrich | G1264 | NA | NA | No | Yes | 9.61E-01 | NA | Protein |
| imipenem | IMI | Water | SigmaAldrich | PHR1796-200MG | NA | NA | No | Yes | Not reached | NA | Cell wall |
| kanamycin | KAN | Water | VWR | 408 | NA | NA | No | Yes | 7.09E-01 | NA | Protein |
| levofloxacin | LEV | DMSO | SigmaAldrich | 28266 | NA | NA | No | Yes | 7.19E-01 | NA | DNA |
| meropenem | MER | DMSO | SigmaAldrich | 1392454 | NA | NA | No | Yes | Not reached | NA | Cell wall |
| mitomycin | MIT | DMSO | SigmaAldrich | Y0000378 | NA | NA | No | Yes | 5.79E-01 | NA | DNA |

| | | | | | | | | | | | |
|---|---|---|---|---|---|---|---|---|---|---|---|
| nigericin | NIG | MeOH | SigmaAldrich | N7143-10MG | NA | NA | No | Yes | 1.23E-01 | NA | Respiration |
| ofloxacin | OFL | NaOH | SigmaAldrich | 8757 | NA | NA | No | Yes | 2.29E-01 | NA | DNA |
| quabodepistat | QBS | DMSO | Otsuka | Gift | NA | NA | No | Yes | 1.66E-01 | NA | Cell wall |
| pseudouridimycin | PSE | DMSO | Fisher Scientific | AG-CN2-0316-M005 | NA | NA | No | Yes | Not reached | NA | RNA |
| SQ109 | SQ1 | DMSO | Fisher Scientific | 50-186-7024 | NA | NA | No | Yes | 5.21E-01 | NA | Cell wall |
| streptomycin | STR | Water | SigmaAldrich | S6501 | NA | NA | No | Yes | 3.28E-01 | NA | Protein |
| tetracycline | TET | DMSO | SigmaAldrich | 87128 | NA | NA | No | Yes | 1.50E+01 | NA | Protein |
| thioridazine | THI | DMSO | Enzo life sciences | BML NS835-0005 | NA | NA | No | Yes | Not reached | NA | Respiration |
| orlistat | THL | DMSO | Enzo life sciences | ALX-350-152-M250 | NA | NA | No | Yes | 2.03E+00 | NA | Cell wall |
| vancomycin | VAN | Water | SigmaAldrich | V0045000 | NA | NA | No | Yes | Not reached | NA | Cell wall |
| verapamil | VER | Water | Enzo life sciences | ALX-55--306-G001 | NA | NA | No | Yes | Not reached | NA | Respiration |
| viomycin | VIO | Water | Fisher Scientific | 37-871-0 | NA | NA | No | Yes | 1.18E+00 | NA | Protein |
| bedaquiline + isoniazid | CB1 | NA | NA | NA | NA | NA | No | Yes | NA | NA | NA |
| bedaquiline + pretomanid | CB2 | NA | NA | NA | NA | NA | No | Yes | NA | NA | NA |
| rifampicin + ethambutol | CB3 | NA | NA | NA | NA | NA | No | Yes | NA | NA | NA |
| moxifloxacin + pretomanid | CB4 | NA | NA | NA | NA | NA | No | Yes | NA | NA | NA |
| rifampicin + sq109 | CB5 | NA | NA | NA | NA | NA | No | Yes | NA | NA | NA |

Supplementary Table 1**. Compounds used in this study.**

| **Morphological Profile Reconstruction Weight** | **RNA-seq Reconstruction Weight** | **Drug Phenotype Classifier Weight** | **Modality Classifier Weight** | **KL-Divergence Weight** | **Epoch** | **# of Latent Dimensions** | **Batch Size** |
|---|---|---|---|---|---|---|---|
| 1 | 1 | 1000 | 1 | 0.1 | 500 | 50 | 16 |

Supplementary Table 2**. DECIPHAER model hyperparameters used for the integration of morphological and RNA-seq profiles of Mtb drug response.**

| **Model** | **Input size** | **Output size** | **# of hidden layers** | **# nodes per layer** | **Activation functions** |
|---|---|---|---|---|---|

| | | | | | |
|---|---|---|---|---|---|
| RNA-seq encoder | 3473 | 50 | 4 | 1024 | ReLu with batch normalization |
| morphological profile encoder | 78 | 50 | 4 | 1024 | ReLu with batch normalization |
| RNA-seq decoder | 50 | 3473 | 4 | 1024 | ReLu with batch normalization |
| morphological profile decoder | 50 | 78 | 4 | 1024 | ReLu with batch normalization |
| drug phenotype classifier | 50 | 11 | 0 | 0 | Linear |
| modality discriminator | 50 | 1 | 3 | 1024 | ReLu |

Supplementary Table 3**. DECIPHAER model architecture used for the integration of morphological and RNA-seq profiles of Mtb drug response.**

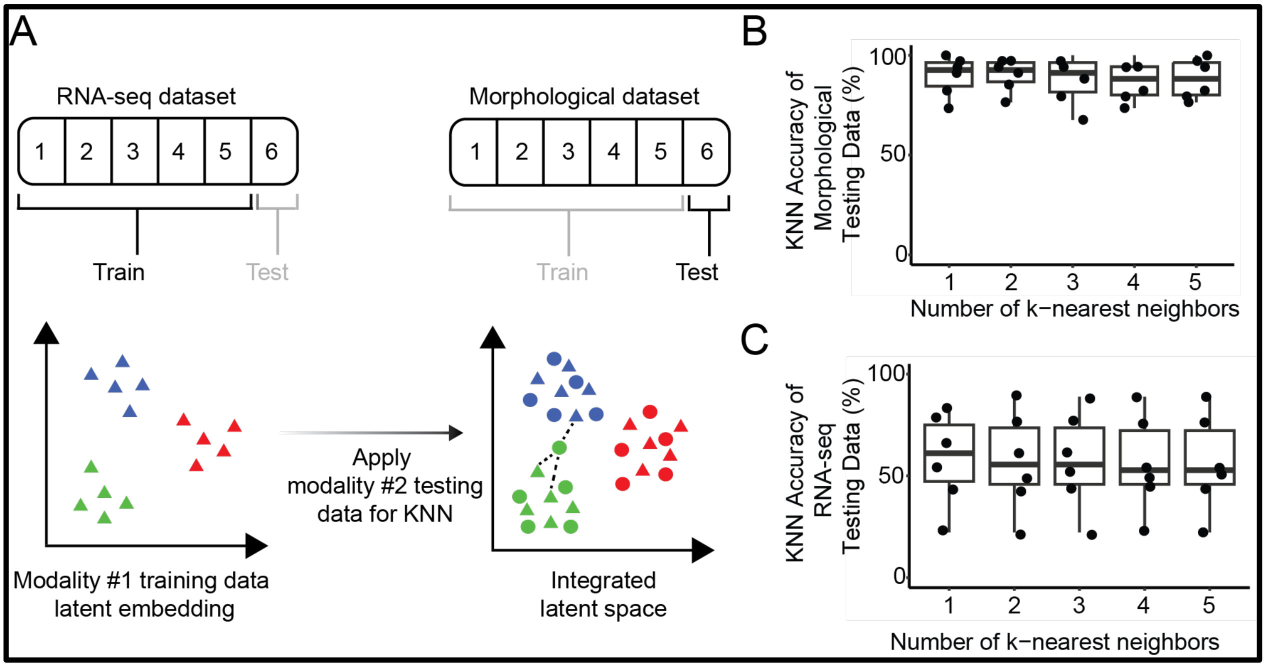


Supplementary Figure 2. **DECIPHAER latent space validation to ensure consistent alignment of morphological and transcriptional profiles.**

(A) Approach for quantifying the degree of cross-modal integration in the latent space. Networks are trained with 6-fold cross-validation. Testing splits of morphological profiling data are applied onto the latent space and the nearest RNA-seq neighbors are identified. If the nearest RNA-seq neighbors are of the same drug phenotype cluster as the morphological testing data, then the KNN accuracy is high.

(B, C) Boxplots displaying kNN accuracy scores (y-axis) across 6-fold cross-validation splits over the number of nearest neighbors considered (x-axis) for the morphological profiling (B) and RNA-seq datasets (C). Each point indicates the testing accuracy for one of the cross-validation splits.

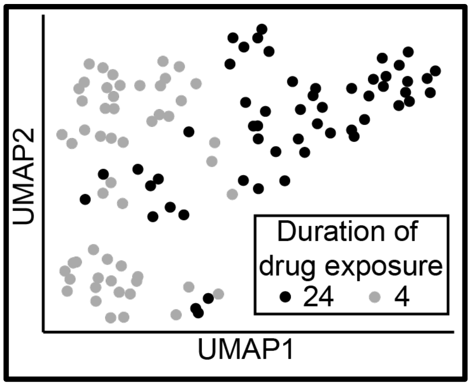


Supplementary Figure 3. **The duration of drug treatment greatly influences gene expression levels in Mtb.** UMAP plot of the DECIPHAER training-set drug treatment RNA-seq data. Each point represents a transcriptomic profile from a biological replicate of Mtb culture treated with a drug for 4- (gray) or 24- (black) hours.

| Latent Embedding F-Test Statistic (Training Data) | Latent Embedding F-Test Statistic (Testing Data) | Morphological Profile Reconstruction Weight | RNA-seq Reconstruction Weight | Drug Phenotype Classifier Weight | Modality Classifier Weight | KL-Divergence Weight | Epoch | # of Latent Dimensions | Batch Size |
|---|---|---|---|---|---|---|---|---|---|
| 41.81 | 17.78 | 1 | 1 | 700 | 1 | 0.1 | 501 | 50 | 16 |
| 124.97 | 45.84 | 1 | 10 | 1000 | 1 | 0.001 | 501 | 50 | 16 |
| 20.22 | 9.59 | 1 | 1 | 500 | 1 | 0.001 | 501 | 50 | 16 |
| 5.31 | 3.42 | 1 | 1 | 1000 | 10 | 0.001 | 601 | 30 | 16 |
| 44.81 | 19.15 | 1 | 10 | 1000 | 1 | 0.01 | 601 | 50 | 16 |
| 8.16 | 3.51 | 1 | 1 | 900 | 1 | 0.001 | 601 | 90 | 16 |
| 104.46 | 36.86 | 1 | 1 | 1000 | 1 | 1.00E-04 | 701 | 50 | 16 |
| 13.02 | 3.12 | 1 | 1 | 2000 | 100 | 1 | 701 | 50 | 16 |

Supplementary Table 4**. F-test statistic values (Methods) of the latent embedded multi-modal training and testing data for a sample of DECIPHAER hyperparameters.**

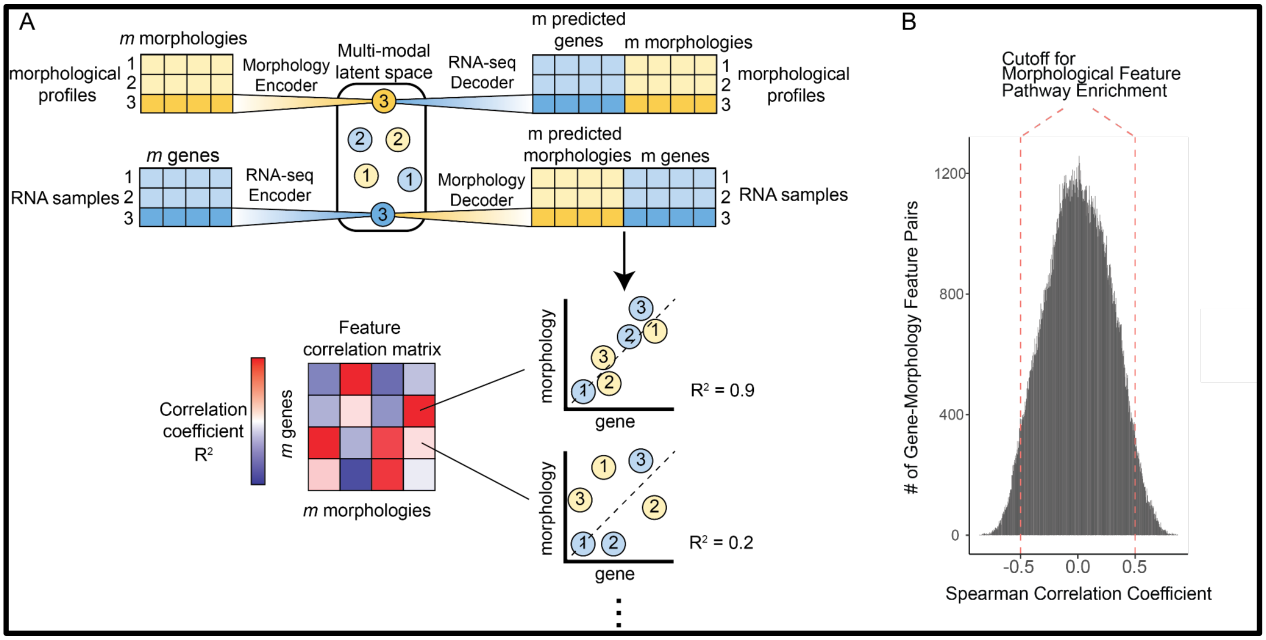


Supplementary Figure 4. **Methodology for using DECIPHAER to investigate cross-modal feature relationships.**

(A) Schematic for DECIPHAER-paired data correlation analysis. After DECIPHAER training, multi-modal latent space data points can be decoded to both RNA-seq and morphological feature values, generating paired measurements (top-right). These data are then used to calculate correlation coefficients between all pairs of morphological and gene expression features (bottom-right) and build a correlation matrix (bottom-left).

(B) Histogram of the Spearman's rank correlation coefficients from all ~6.3 million cross-modal feature pairs. The dashed lines indicate the cutoff used for biological pathway enrichment analysis (> 0.5 or < -0.5).

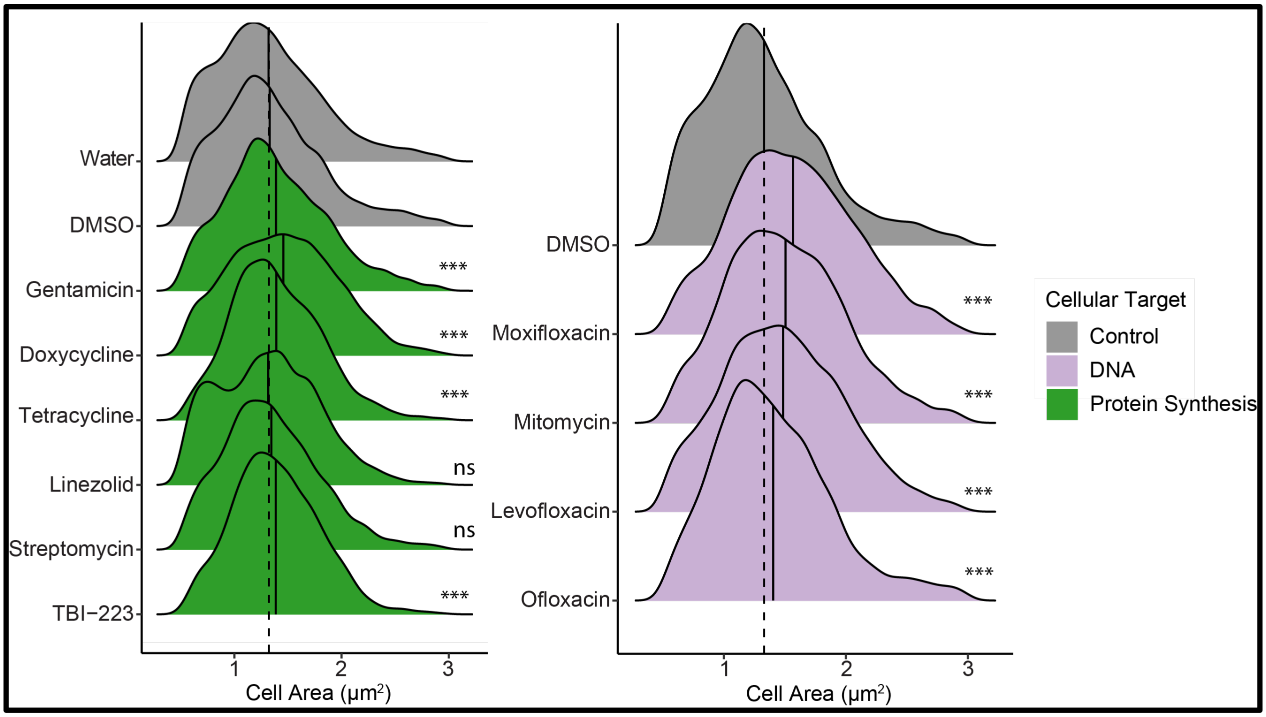


Supplementary Figure 5. **Cell size morphological features are influenced by perturbations to protein and DNA synthesis.** Cell area density plots of single-cell Mtb treated with high doses (10x IC50) of protein and DNA synthesis inhibitors. The solid line indicates the median. The dashed line shows the median of untreated cells (One-way ANOVA with Dunnett's post-hoc test against DMSO control: ***$p < 0.001$, **$p < 0.01$, *$p < 0.05$; ns, $p > 0.05$). Each treatment group had at least a total of 4,509 cells from 3 separate experiments.

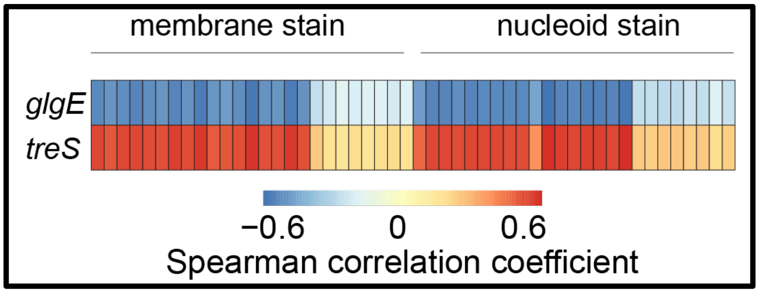


Supplementary Figure 6. **Cell staining features correlate with sugar metabolism gene expression.** Heatmap of the Spearman's correlation coefficients between all morphological profile staining features (FM464 and SYTO24) and *treS* and *glgE* gene expression. Coefficients were calculated using DECIPHAER-paired data (Methods).

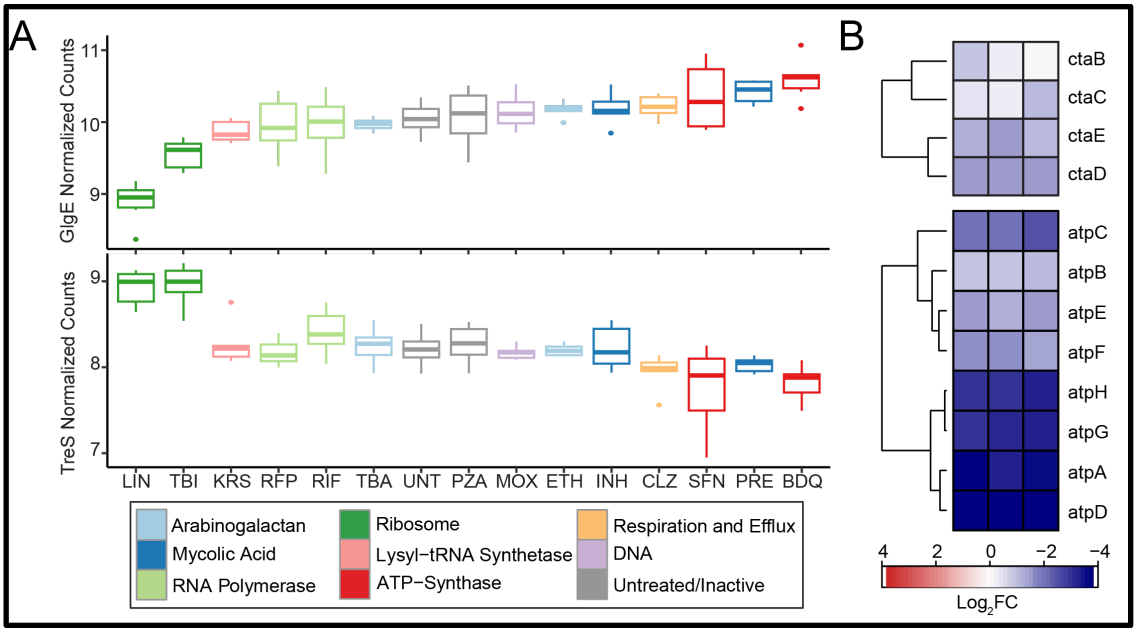


Supplementary Figure 7. **Mtb treS and glgE pathway gene expression response to TB drug treatments.** (A) Boxplots of the variance stabilized transformed (VST) *glgE* (top) and *treS* (B) transcript read counts from 4 and 24-hour treatment time-points (n = 3, with technical duplicate for untreated). (B) Respiration metabolism gene expression response of Mtb to linezolid. Fold change in VST transformed read counts of cytochrome c oxidase (top) and ATP-synthase (bottom) after 24 hours of linezolid treatment (n=3). Fold change is relative to untreated.

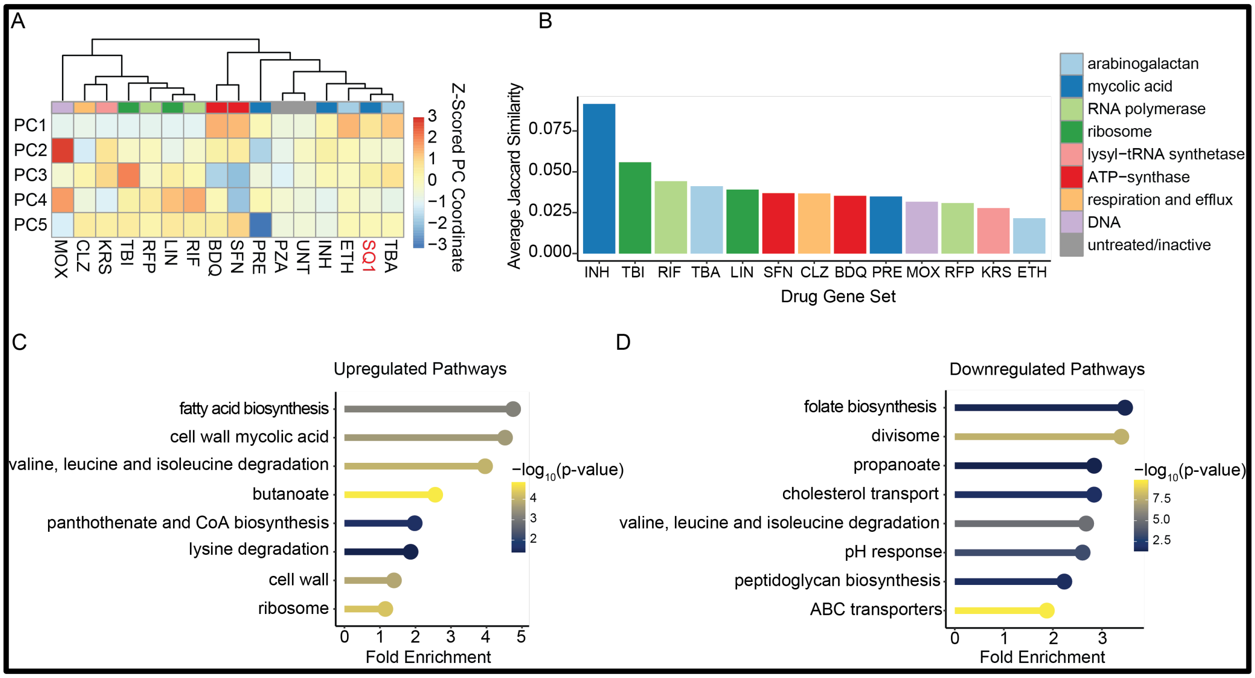


Supplementary Figure 8. **Single-modal analysis demonstrates SQ109's cell wall impact.** (A) Hierarchical clustering on the median morphological profiles of SQ109 and all training drugs projected on the first five principal components. rows are z-scored to demonstrate relative

differences between drug treatments. The dendrogram was computed using Manhattan distances and Ward2 clustering.
(B) Average Jaccard similarity coefficient between predicted RNA-seq of Mtb treated with SQ109 and all training drugs' translated RNA-seq data. Annotation colors in (A) and (B) indicate the cellular target of the drug treatment.
Up- (C) and down- (D) regulated pathway enrichments of SQ109 RNA-seq data predicted from morphological data. The differential expression testing for pathway enrichment compared SQ109 against the training-set translated-RNA-seq through Wilcoxon rank-sum tests. The color of the bubble plot indicates the highest -$\log_{10}$(p-value) among the differentially expressed genes in the pathway and the x-axis indicates the fold enrichment of the pathway (PathFindR).

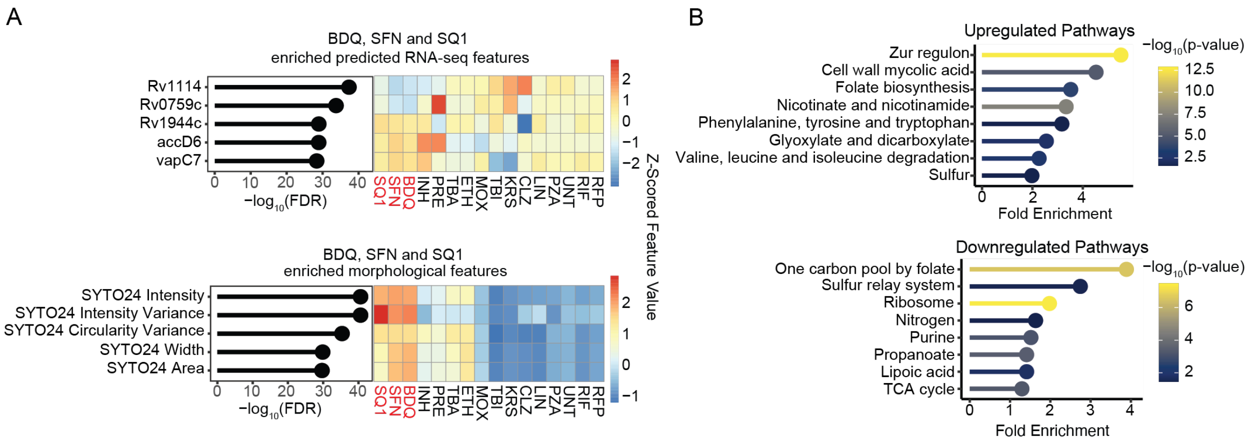


Supplementary Figure 9. **Morphological- and predicted RNA-seq features discriminating SQ109, SFN, and BDQ.**
(A) Clustered heatmaps of the top 5 most enriched predicted RNA-seq features (top) and morphological features (bottom) in Mtb treated with BDQ, SFN, and SQ109. The left side indicates the feature's enrichment level in Welch's t-tests (top) and Wilcoxon rank-sum tests (bottom) with Benjamini-Hochberg (FDR) correction comparing BDQ, SFN, and SQ109 against all other compounds and untreated. The heatmap displays the median feature value (z-scored) for each drug.
(B) Pathway enrichment of transcriptional data translated from morphological profiles of Mtb treated with BDQ, SFN, and SQ109. The differential expression testing for pathway enrichment compared BDQ, SFN, and SQ109 against all training-set translated RNA-seq through Wilcoxon rank-sum tests. The color of the bubble plot indicates the highest -$\log_{10}$(p-value) among the differentially expressed genes in the pathway and the x-axis indicates the fold enrichment of the pathway (PathFindR).

| **Drug** | **Most similar clade drug 1** | **Most similar clade drug 2** | **Most similar clade drug 3** | **Expected broad cellular target** | **Broad cellular target of drug 1 in clade** | **Broad cellular target of drug 2 in clade** | **Broad cellular target of drug 3 in clade** | **Testing drug shares clade with same broad cellular target** |
|---|---|---|---|---|---|---|---|---|

| | | | | | | | | |
|---|---|---|---|---|---|---|---|---|
| actinomycin D | LIN | RIF | RFP | RNA | Protein | RNA | RNA | In Clade |
| amikacin | LIN | RIF | RFP | Protein | Protein | RNA | RNA | In Clade |
| ampicillin | INH | NA | NA | Cell wall | Cell wall | | | In Clade |
| chloramphenicol | TBI | NA | NA | Protein | Protein | | | In Clade |
| capreomycin | CLZ | NA | NA | Protein | Respiration | | | Not in Clade |
| CCCP | LIN | RIF | RFP | Respiration | Protein | RNA | RNA | Not in Clade |
| cerulenin | PRE | NA | NA | Cell wall | Cell wall | | | In Clade |
| ciprofloxacin | MOX | NA | NA | DNA | DNA | | | In Clade |
| clarithromycin | LIN | RIF | RFP | Protein | Protein | RNA | RNA | In Clade |
| cefotaxime | ETH | TBA | NA | Cell wall | Cell Wall | | | In Clade |
| cycloserine | CLZ | NA | NA | Cell wall | Respiration | | | Not in Clade |
| daunorubicin | MOX | NA | NA | DNA | DNA | | | In Clade |
| delamanid | PRE | NA | NA | Cell wall | Cell wall | | | In Clade |
| doxycycline | LIN | RIF | RFP | Protein | Protein | RNA | RNA | In Clade |
| ethionamide | ETH | TBA | NA | Cell wall | Cell wall | | | In Clade |
| fidaxomicin | LIN | RIF | RFP | RNA | Protein | RNA | RNA | In Clade |
| gentamicin | DD1 | NA | NA | Protein | Protein | | | In Clade |
| kanamycin | MOX | NA | NA | Protein | DNA | | | Not in Clade |
| levofloxacin | MOX | NA | NA | DNA | DNA | | | In Clade |
| mitomycin | MOX | NA | NA | DNA | DNA | | | In Clade |
| nigericin | CLZ | NA | NA | Respiration | Respiration | | | In Clade |
| ofloxacin | DD1 | NA | NA | DNA | Protein | | | Not in Clade |
| quabodepistat | ETH | TBA | NA | Cell wall | Cell wall | | | In Clade |

| | | | | | | | | |
|---|---|---|---|---|---|---|---|---|
| SQ109 | BDQ | NA | NA | Cell wall | Respiration | | | Not in Clade |
| streptomycin | MOX | NA | NA | Protein | DNA | | | Not in Clade |
| tetracycline | MOX | NA | NA | Protein | DNA | | | Not in Clade |
| orlistat | CLZ | NA | NA | Cell wall | Respiration | | | Not in Clade |
| viomycin | DD1 | NA | NA | Protein | Protein | | | In Clade |

Supplementary Table 5. **Cellular target classification through unsupervised clustering of latent embedded morphological profiles of drugs for which we did not measure RNA-seq.** The median values of the latent space features for the testing drugs and all training drugs were hierarchically clustered with Manhattan distances and Ward2 clustering. The neighbors of the testing drug in the most similar clade (max of 3) were determined.

| Drug morphological profile that was translated to RNA-seq | Broad cellular target of translated data | Full name of training drug with most similar transcriptional profile | Broad cellular target of most similar training drug | Jaccard Similarity |
|---|---|---|---|---|
| actinomycin D | RNA | rifampicin | RNA | 0.32 |
| amikacin | Protein | rifampicin | RNA | 0.363 |
| ampicillin | Cell wall | isoniazid | Cell wall | 0.07 |
| chloramphenicol | Protein | krs | Protein | 0.208 |
| capreomycin | Protein | clofazimine | Respiration | 0.095 |
| CCCP | Respiration | clofazimine | Respiration | 0.145 |
| cerulenin | Cell wall | moxifloxacin | DNA | 0.111 |
| ciprofloxacin | DNA | ethambutol | Cell wall | 0.09 |
| clarithromycin | Protein | rifampicin | RNA | 0.123 |
| cefotaxime | Cell wall | pretomanid | Cell wall | 0.107 |
| cycloserine | Cell wall | clofazimine | Respiration | 0.139 |
| daunorubicin | DNA | moxifloxacin | DNA | 0.165 |
| delamanid | Cell wall | pretomanid | Cell wall | 0.053 |

| | | | | |
|---|---|---|---|---|
| doxycycline | Protein | linezolid | Protein | 0.144 |
| ethionamide | Cell wall | ethambutol | Cell wall | 0.48 |
| fidaxomicin | RNA | linezolid | Protein | 0.145 |
| gentamicin | Protein | krs | Protein | 0.112 |
| kanamycin | Protein | krs | Protein | 0.047 |
| levofloxacin | DNA | moxifloxacin | DNA | 0.235 |
| mitomycin | DNA | moxifloxacin | DNA | 0.135 |
| nigericin | Respiration | clofazimine | Respiration | 0.246 |
| ofloxacin | DNA | krs | Protein | 0.122 |
| quabodepistat | Cell wall | tba-7371 | Cell wall | 0.172 |
| sq109 | Cell wall | isoniazid | Cell wall | 0.091 |
| streptomycin | Protein | krs | Protein | 0.063 |
| tetracycline | Protein | moxifloxacin | DNA | 0.129 |
| orlistat | Cell wall | rifampicin | RNA | 0.086 |
| viomycin | Protein | krs | Protein | 0.097 |

Supplementary Table 6. **Cellular target classification of testing drugs through average Jaccard similarity of their predicted transcriptional data.** Differential expression lists were calculated for each drug's predicted RNA-seq profile and compared with those of drugs in the training set. The training drug with the highest average Jaccard similarity coefficient of differential expression (Methods) is reported.

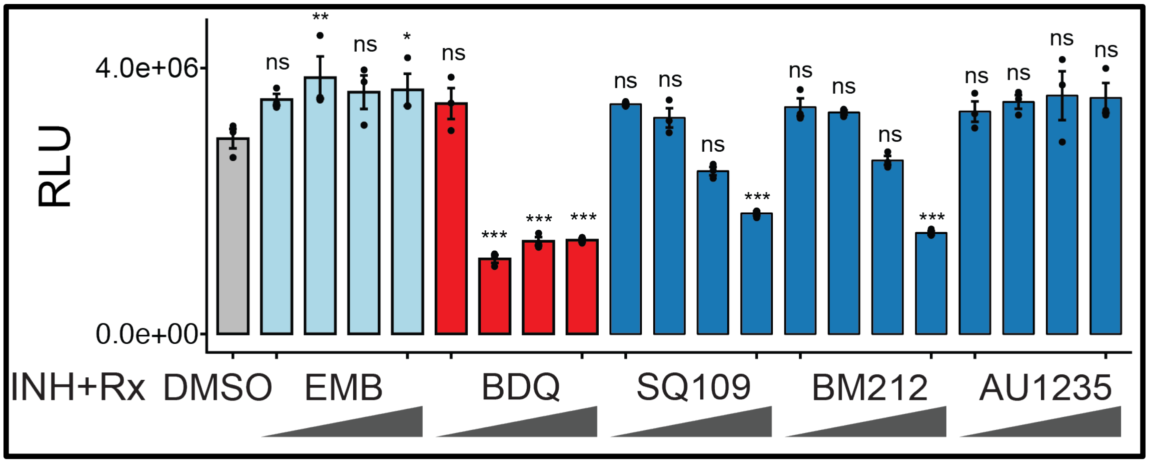


Supplementary Figure 10. **Quantification of ATP content in drug-treated Mtb MmpL3-F644L-overexpression cultures.** The inducible MmpL3-F644L Mtb strain was cultured in the presence of ATc for 4.5 generations to cause overexpression of the mutant protein. Cultures were then treated with DMSO, INH, or INH in combination with four different doses at 0.1x-, 0.75x-, 5x-, and 10x- IC50 of a second drug for 24 hours. The INH treatment dose was held constant at 10x IC50. ATP content was measured with the BacTiter-Glo$^{TM}$ assay. Luminescence relative to culture $OD_{600nm}$ (RLU) is plotted (Barplots depict the mean value and error bars are calculated from the standard error of the mean. One-way ANOVA with Dunnett's post-hoc test against DMSO control: ***$p < 0.001$, **$p < 0.01$, *$p < 0.05$; ns, $p > 0.05$; n = 3).One-way ANOVA with Dunnett's post-hoc test against INH+DMSO control: ***$p < 0.001$, **$p < 0.01$, *$p < 0.05$; ns, $p > 0.05$; n = 3).
